# A *Peronophythora litchii* RXLR effector PlAvh202 enhances plant susceptibility by destabilizing host SAMS

**DOI:** 10.1101/2022.08.22.504805

**Authors:** Peng Li, Wen Li, Xiaofan Zhou, Junjian Situ, Lizhu Xie, Pinggen Xi, Bo Yang, Guanghui Kong, Zide Jiang

## Abstract

Oomycete pathogens can secret hundreds of effectors into plant cells to interfere with plant immune system during infection. Here we identified a cytoplasmic RXLR effector from the most destructive pathogen of litchi, *Peronophythora litchii*, and named it as PlAvh202. PlAvh202 could suppress cell death triggered by INF1 and Avr3a/R3a in *Nicotiana benthamiana*, and was essential for *P. litchii* virulence. In addition, PlAvh202 also suppressed plant immune responses and promoted the susceptibility of *N. benthamiana* to *Phytophthora capsici*. Further research revealed that PlAvh202 could suppress ethylene (ET) production by targeting and destabilizing plant S-adenosyl-L-methionine synthetase (SAMS), a key enzyme in ET biosynthesis pathway, in a 26S proteasome-dependent manner. Transient expression of LcSAMS3 induced ET production and enhanced plant resistance, whereas inhibition of ET biosynthesis promoted *P. litchii* infection, supporting that LcSAMS and ET positively regulate litchi immunity towards *P. litchii*. Overall, these findings highlight that SAMS can be targeted by oomycete RXLR effector to manipulate ET-mediated plant immunity.

## Introduction

Both pathogens and plants have evolved diverse pathways to counter each other for more beneficial growing condition in the long process of co-evolution. To resist the invasion of pathogens, plants have two layers of defense systems. Firstly, some conserved pathogen-associated molecular patterns (PAMPs) can be recognized by plant cell surface pattern-recognition receptors (PRRs) to trigger basal defense response, known as PAMP-triggered immunity (PTI) (Jones and Dangl, 2006; DeFalco and Zipfel, 2021). For instance, bacterial flagellin, elongation factor Tu, and *Phytophthora infestans* elicitin infestin 1 (INF1) are typical PAMPs which are recognized by *Arabidopsis* FLS2, EFR, and potato RLP85, respectively, leading to promoted plant resistance (Gomez-Gomez and Boller, 2000; Zipfel et al., 2006; Du et al., 2015). However, a variety of pathogens can secrete virulence-related molecules, such as effectors, to suppress plant PTI for successful infection. As a response, plants initiate the second layer of defense with the resistance proteins (R proteins) activation. These R proteins can specifically recognize pathogen effectors directly or indirectly to cause a cascade of downstream immune responses; this process is named effector-triggered immunity (ETI) (Yuan et al., 2021). For example, the avirulence protein AVR3a^K80I103^ from *P. infestans* is specifically recognized by potato resistance protein R3a to trigger ETI-mediated cell death (Armstrong et al., 2005). *Pseudomonas syringae* effector AvrRpt2 indirectly actives R protein RPS2 by eliminating *Arabidopsis* protein RIN4, which induces a series of ETI responses (Axtell and Staskawicz, 2003). Although PTI and ETI are initiated by diverse mechanisms, they both can result in reactive oxygen species (ROS) accumulation and hypersensitive response (HR), a form of programmed cell death (PCD) that can restrict pathogens proliferation (Adachi et al., 2015; Zebell and Dong, 2015; Li et al., 2019).

Oomycetes comprise numerous destructive plant pathogens such as *P. infestans*, *Phytophthora sojae*, and *Peronophythora litchii*, which are great threats to economic crops. Thus, understanding the pathogenic mechanism is significant to restrict the invasion of pathogens for higher crop yields. Oomycetes secret numerous effectors into plant cells to interfere with plant immune system, among which RXLR effectors have a significant contribution for pathogens virulence (Jiang et al., 2008). The RXLR effectors exhibit sequence polymorphisms in general, but they possess two highly conserved motifs, termed RXLR (Arg-X-Leu-Arg; X is any amino acid) and dEER (Asp-Glu-Glu-Arg; Asp is less conserved than other three amino acids), following the N-terminal signal peptide (SP) (Wawra et al., 2012). Previous reports have shown that the RXLR motif is involved in plant cell entry (Whisson et al., 2007; Kale et al., 2010; Kale and Tyler, 2011). RXLR effectors have various functions in plant-pathogen interaction, including both eliciting and suppressing cell death, under different circumstance. For example, AVR2 and Pi17316 from *P. infestans* could suppress INF1-triggered cell death (ICD) in *N. benthamiana* depending on CHL1 and VIK, respectively (Turnbull et al., 2017; Murphy et al., 2018). Avr1b of *P. sojae* could suppress BAX-triggered cell death (Dou et al., 2008). On the other hand, *P. sojae* Avh241 and *P. capsici* RXLR207 could trigger cell death in *N. benthamiana* (Yu et al., 2012; Li et al., 2019). Large-scale screening assays of *P. sojae, Hyaloperonospora arabidopsidis*, and *Plasmopara viticola* suggest that most RXLR effectors can suppress cell death triggered by several elicitors in *N. benthamiana*, whereas a few RXLR effectors trigger cell death (Fabro et al., 2011; Wang et al., 2011; Liu et al., 2018). These data support a hypothesis that most RXLR effectors secreted from pathogens mainly aim at suppressing host immune system for infection. Thus, the study that effector-mediated PTI or ETI suppression is very valuable to reveal the mechanism of plant-pathogen interactions.

Salicylic acid (SA), jasmonic acid (JA), and ET are three major phytohormones in plant defense responses (Casteel et al., 2015). In plants, the biosynthesis of ET begins with the conversion of L-methionine (L-Met) and ATP into S-adenosyl-L-methionine (AdoMet, SAM), catalyzed by S-adenosyl-L-methionine synthetase (SAMS) [EC 2.5.1.6]. SAM is then converted to 1-aminocyclopropane-1-carboxylic acid (ACC) and methylthioadenosine (MTA), by 1-aminocyclopropane-1-carboxylate synthase (ACS). ACC can be oxidized by ACC oxidases (ACO) to produce ET (Park et al., 2021). Because SAM acts as a precursor of ET and polyamines, SAMS is important for ET production. The SAMS enzyme can be induced by biotic and abiotic stress and participate in the methylation of DNA or histone to regulate plant development (Li et al., 2011). In addition, SAMS is also involved in the regulation of plant resistance. For example, *Cotton Leaf Curl Multan virus* (CLCuMuV) C4 protein reduces the SAMS enzymatic activity to repress transcriptional gene silencing (TGS) and post-transcriptional gene silencing (PTGS), which can enhance CLCuMuV infection in plants (Ismayil et al., 2018). Given that SAMS plays a role in plant development and resistance, it is significant to explore the effect of SAMS on plant-pathogen interactions.

Previous study has predicted 245 RXLR effectors in *P. litchii* (Ye et al., 2016), whereas their function in immune suppression is not fully understood. In this study, we found four *P. litchii* RXLR effectors, PlAvh42, PlAvh202, PlAvh208, and PlAvh222, that could suppress ICD, based on a large-scale screening in *N. benthamiana*. Among them, PlAvh202 had a strong ability to suppress cell death triggered by INF1 or Avr3a/R3a, and was required for *P. litchii* virulence. Therefore, we focused our investigation on PlAvh202 and found that PlAvh202 suppressed plant innate immune responses, including suppressing immune maker gene expression and ROS accumulation. In addition, we found that PlAvh202 could interact with *N. benthamiana* NbSAMS proteins as well as the orthologous SAMS in litchi, designated as LcSAMS, via its core virulence region IR2 (internal repeat). Furthermore, PlAvh202 could not promote *P. capsici* infection in the absence of IR2 or in *NbSAMS*s-silenced *N. benthamiana*, suggesting that PlAvh202 exerts its virulence by targeting SAMS. In summary, our results reveal a mechanism that an RXLR effector PlAvh202 can destabilize plant SAMS via 26S proteasome to suppress ET-mediated plant immunity.

## Results

### PlAvh202 can suppress leaf cell death of *N. benthamiana* triggered by INF1 or Avr3a/R3a

Programed cell death (PCD) or hypersensitive response (HR) is a significant characteristic of PTI and ETI, and a number of effectors have been confirmed to suppress plant PCD or HR to disrupt PTI or ETI (Abramovitch et al., 2003; Liu et al., 2011; Dutra et al., 2020). In this study, each of 48 RXLR effectors of *P. litchii* without signal peptide (SP) was transiently expressed in *N. benthamiana* leaves by *Agrobacterium tumefaciens* infiltration to test whether they could suppress ICD (Table S1). INF1 was transiently expressed in the same infiltration region after 24 hours. Eventually, we found that INF1 could not trigger cell death in *N. benthamiana* leaves when PlAvh42, PlAvh202, PlAvh208 or PlAvh222 was expressed individually. Meanwhile, the expression of RFP control in leaves did not suppress the cell death triggered by INF1 (Figure 1A). This result suggested that these four effectors, namely PlAvh42, PlAvh202, PlAvh208, and PlAvh222, could suppress ICD.

**Figure 1.**
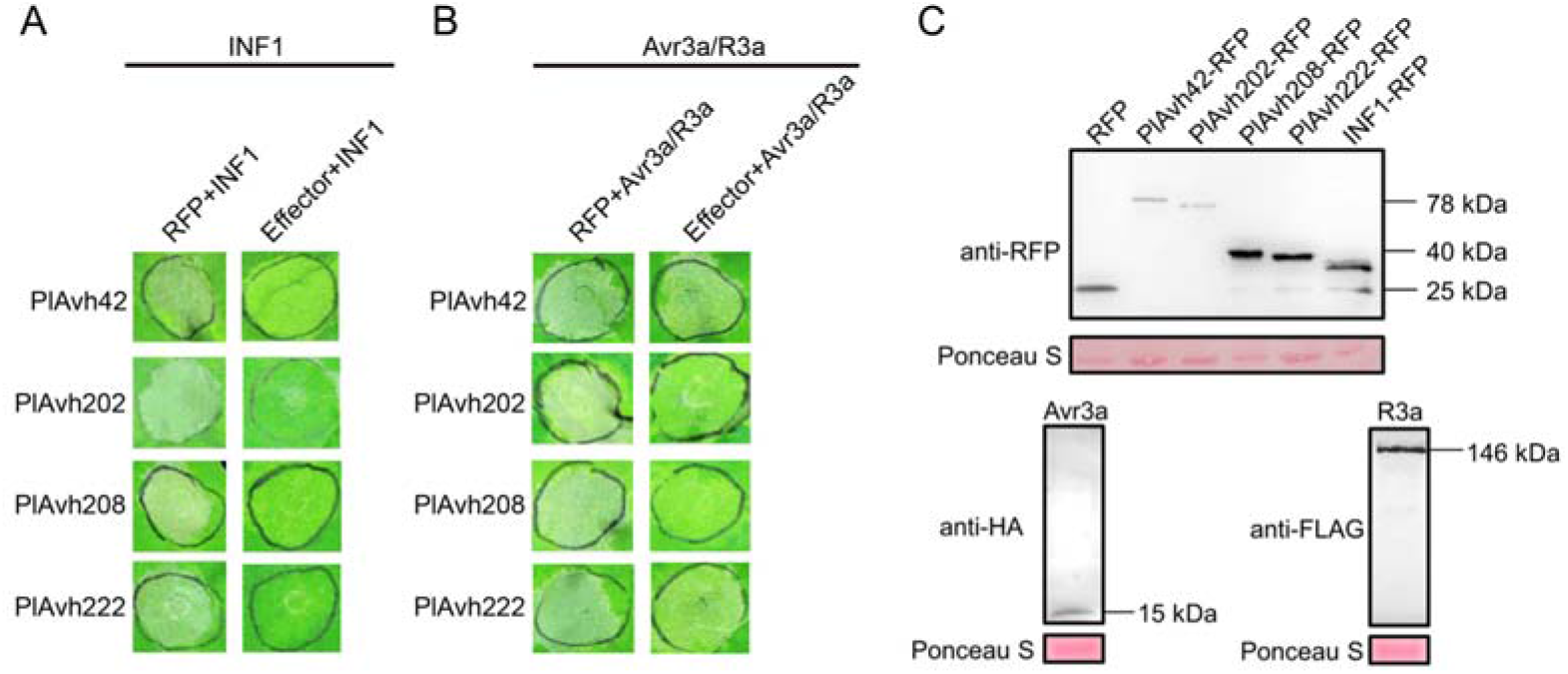
*Peronophythora litchii* RXLR effectors suppress cell death triggered by INF1 or Avr3a/R3a in *N. benthamiana*. **(A, B)** Partial tissue responses to INF1 **(A)** or Avr3a/R3a **(B)** in the presence of *P. litchii* RXLR effectors. PlAvh202, PlAvh208, and PlAvh222 suppressed cell death triggered by INF1 or Avr3a/R3a; PlAvh42 only suppressed cell death triggered by INF1. Effector-encoding genes were transiently expressed in *N. benthamiana* leaves by agroinfiltration, then *INF1* or *Avr3a/R3a* was transiently expressed in the indicated regions 24 h later. RFP was used as control. Photographs were taken at 3 days post-agroinfiltration (dpa). **(C)** Immunoblot analysis. All proteins tested in the assay were confirmed using western blot, and total protein was stained by Ponceau S.

Next, we examined whether these four effectors were able to suppress cell death triggered by Avr3a/R3a in *N. benthamiana* leaves. The experiment result showed that three out of these four RXLR effectors, PlAvh202, PlAvh208 and PlAvh222 could also suppress leaf cell death triggered by Avr3a/R3a, whereas PlAvh42 could not (Figure 1B). Protein expression of four tested effectors, INF1, and Avr3a/R3a were verified by western blot (Figure 1C). Overall, these results indicated that PlAvh202, PlAvh208, and PlAvh222 could suppress plant cell death mediated by INF1 or Avr3a/R3a, while PlAvh42 only suppressed ICD. Given that PlAvh202 showed the strongest inhibitory effect on *N. benthamiana* leaves cell death among the four tested effectors, we selected PlAvh202 for further investigation.

### PlAvh202 is critical for *P. litchii* virulence

To investigate the potential pathogenicity and virulence of *PlAvh202*, we examined the expression pattern of *PlAvh202* using reverse transcription-quantitative polymerase chain reaction (RT-qPCR). The result showed that *PlAvh202* was substantially up-regulated in zoospores and at 1.5, 3, 6 hours post-inoculation (hpi) compared with the mycelial stage, and the highest expression level occurred in the zoospore stage (more than 1800-fold) (Figure S1). This result suggested that *PlAvh202* is highly expressed during early infection stages of *P. litchii*, and indicated that *PlAvh202* may contribute to the virulence of *P. litchii*.

To further explore the contribution of *PlAvh202* to *P. litchii* virulence, we deleted the *PlAvh202* gene in *P. litchii* by CRISPR/Cas9 technology (Fang et al., 2017) (Figure 2A, S2A). Three transformants (T7, T55, T95) were verified as successful *PlAvh202* deletion mutants, based on PCR amplification and Sanger sequencing (Figure 2B, S2B). A transformant that failed to delete *PlAvh202* was selected as control (CK). We next evaluated the virulence of these three *PlAvh202* deletion mutants by inoculating their zoospores on litchi leaves, and the result showed that loss of *PlAvh202* gene caused reduced virulence, as smaller lesions were formed, compared to those caused by wild-type (WT) or CK inoculation (Figure 2C, D). This result suggested that *PlAvh202* was required for the full virulence of *P. litchii*.

**Figure 2.**
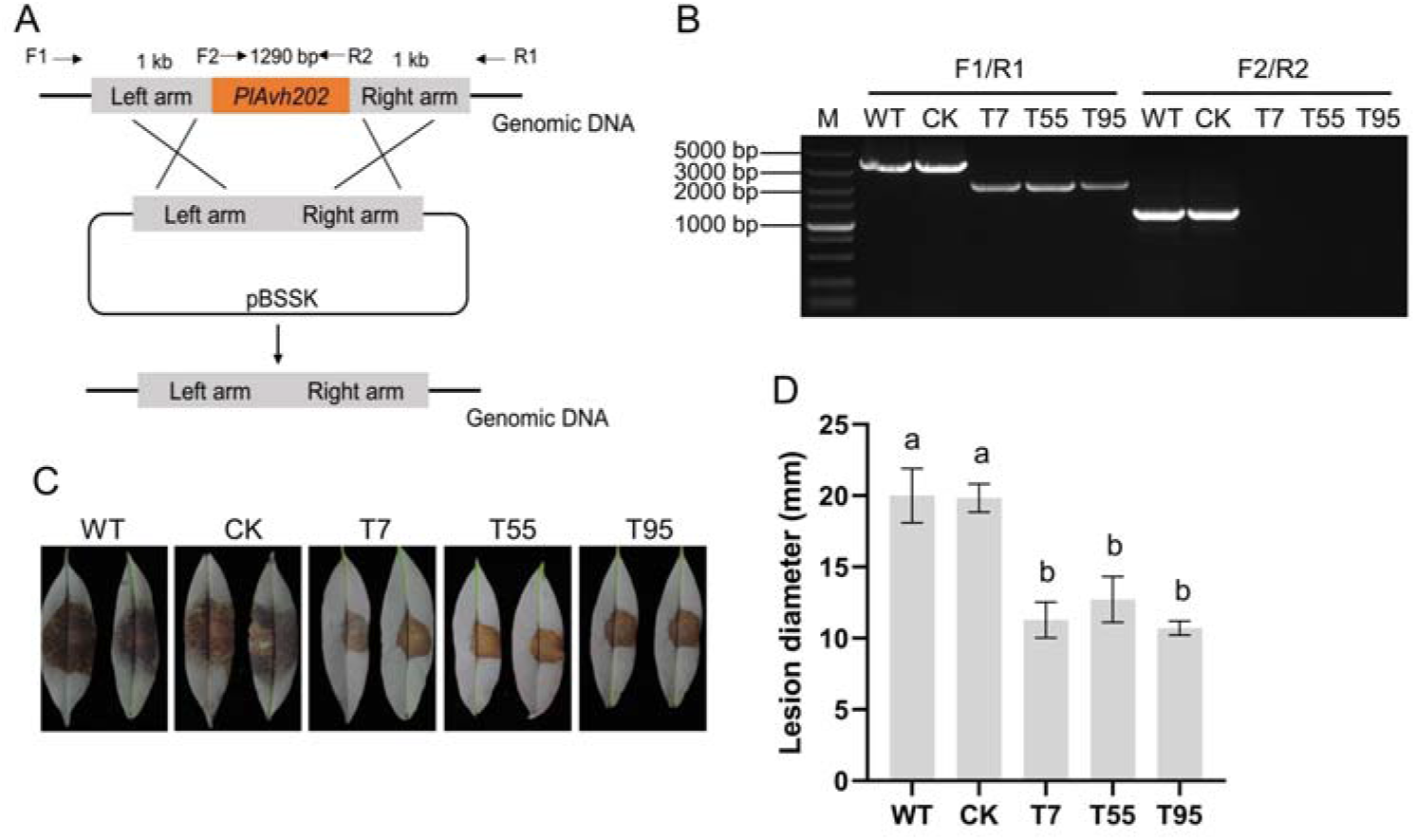
Deletion of *PlAvh202* impairs the virulence of *P. litchii*. **(A)** Schematic diagram of *PlAvh202* deletion using CRISPR/Cas9. The primer pairs F1/R1 and F2/R2 used for *PlAvh202* mutants screening are indicated by the horizontal arrows. **(B)** PCR analysis of *PlAvh202* mutants. Genomic DNA of WT, CK, T7, T55, and T95 was used for PCR assays by primer pairs F1/R1 and F2/R2, respectively. **(C)** Virulence assays of *PlAvh202* mutants on litchi leaves. 100 zoospores of WT, CK, T7, T55 and T95 were inoculated on the center of litchi leaves. Photographs were taken at 48 h post-inoculation (hpi). **(D)** Lesion diameter of *P. litchii* infection at 48 hpi. Different letters represent significant differences using the one-way ANOVA test followed by Tukey’s honestly significant difference (HSD) test (*P*< 0.05). Data are the means ± SD of three independent biological replicates (n≥6 leaves).

In addition, we also measured mycelial growth rates and found no statistically significant difference among WT, CK, T7, T55, and T95 in their colony diameters (Figure S2C, 2D), indicating that *PlAvh202* deletion did not impact the mycelial growth of *P. litchii*.

### PlAvh202 suppresses *N. benthamiana* immune responses induced by INF1 and promoted pathogen infection

Because of the stable ability of PlAvh202 to suppress PCD induced by INF1 in *N. benthamiana* leaves, we further explored its effect on the plant immune system. We evaluated the relative expression levels of maker genes related to SA, JA, ET, and ROS accumulation, including *PR1/2* (pathogenesis-related protein 1/2), *LOX* (lipoxygenase), *PDF1.2* (Plant defensin1.2), *RbohA*, and *RbohB* (respiratory burst oxidase homologs) (Yoshioka et al., 2003; Pieterse et al., 2012; Yang et al., 2019; Ding and Ding, 2020), using RT-qPCR. The result showed that the transient expression of PlAvh202 led to the reduced expression levels of *NbPR1*, *NbPR2*, *NbLOX*, *NbRbohA*, and *NbRbohB* at 24, 36, and 48 hpi. Although ET/JA pathway-related maker gene *NbPDF1.2* was up-regulated at 24 hpi, its expression level was also down-regulated at 36 and 48 hpi (Figure 3A). Together, these maker genes were suppressed by PlAvh202 at 36 and 48 hpi. Given that *NbRbohA* and *NbRbohB* are required for ROS accumulation in *N. benthamiana* to resist pathogens, it is speculated that PlAvh202 might reduce ROS production of *N. benthamiana*. To validate this hypothesis, we tested the ROS accumulation of *N. benthamiana* leaves using DAB staining after expressing PlAvh202 and INF1 successively. The brown area where PlAvh202 was transiently expressed was significantly smaller than that of RFP (Figure 3B), suggesting PlAvh202 could attenuate INF1-triggered ROS accumulation in *N. benthamiana*. Thus, we conclude that PlAvh202 can suppress *N. benthamiana* immune responses induced by INF1.

**Figure 3.**
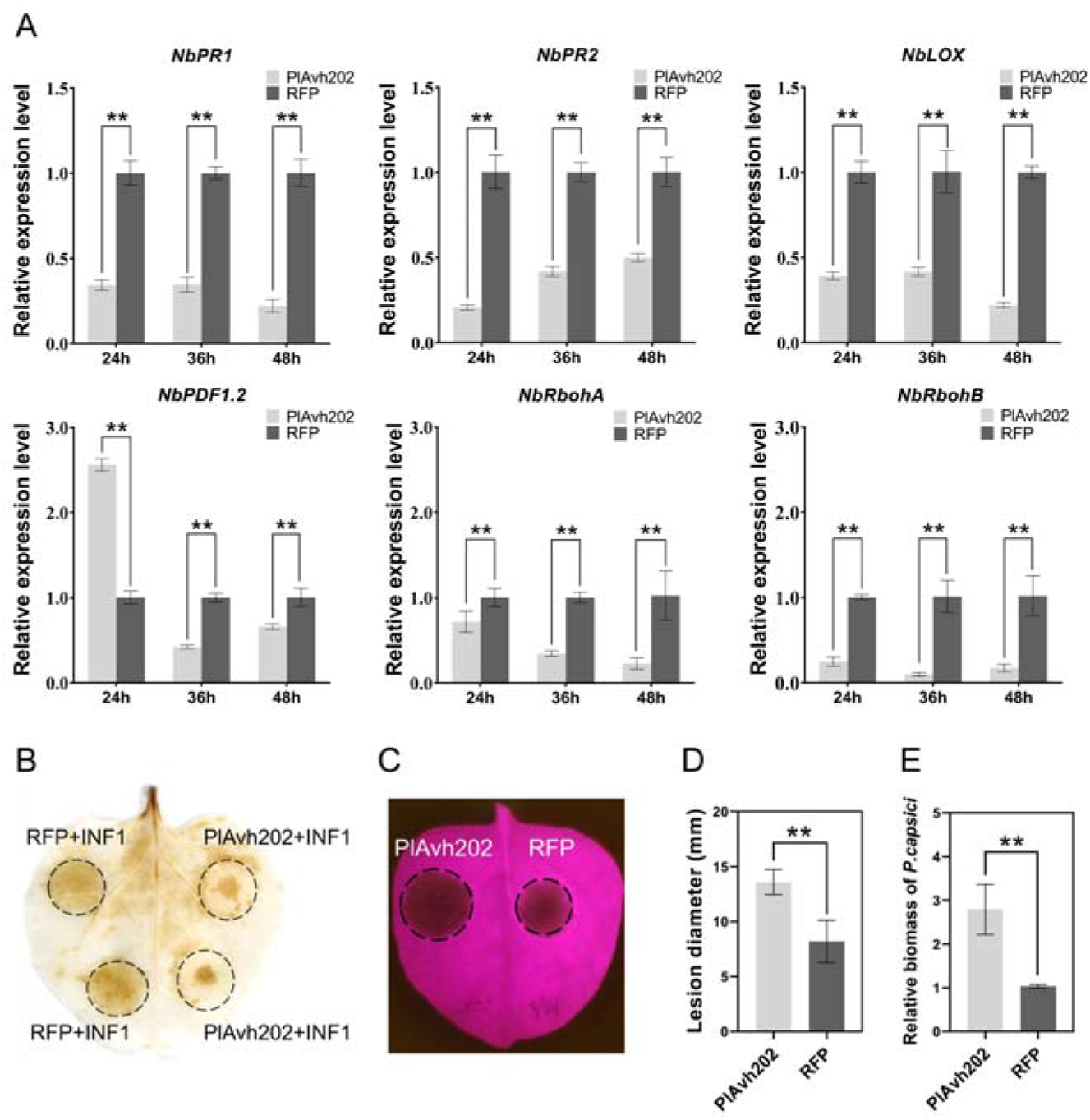
PlAvh202 suppresses INF1-triggered plant immune responses. **(A)** Expression profile analysis of immune maker genes. Relative expression levels of *NbPR1*, *NbPR2*, *NbLOX*, *NbPDF1.2*, *NbRbohA*, and *NbRbohB* induced by INF1 in the presence of PlAvh202 or RFP control were measured using RT-qPCR. The total RNA of *N. benthamiana* leaves was extracted at 24, 36 and 48 hpi. Data are the means ± SD of three independent biological replicates (Each sample of every experiment were from 3 leaves at each timepoint), asterisks represent significant differences (\*\**P* < 0.01, Student’s *t* test). *NbEF1a* was used as the internal reference. **(B)** ROS accumulation in *N. benthamiana*. The leaves were stained by 1 mg/mL DAB at 36 h after agroinfiltration carrying *PlAvh202*, *INF1*, or *RFP*. The black circles indicate the inoculation region. **(C)** PlAvh202 promoted *P. capsici* infection in *N. benthamiana* leaves. *N. benthamiana* leaves were infiltrated with agrobacterium carring *RFP* or *PlAvh202*, then the infiltrated leaves were inoculated with *P. capsici* at 24 h after infiltration. Photographs were taken under UV light at 36 hpi. Black circles indicate lesion areas. **(D, E)** Lesion diameter **(D)** and biomass **(E)** of *P. capsici* in *N. benthamiana* leaves expressing PlAvh202 or RFP. Data are the means ± SD of three independent biological replicates (n≥3), asterisks represent significant differences (\*\**P* < 0.01, Student’s *t* test). *PcActin* and *NbEF1α* were used for qPCR to analyze the biomass of *P. capsici*.

In order to demonstrate whether overexpression of PhAvh202 increases plant susceptibility to pathogens, we inoculated *P. capsici* on both sides of *N. benthamiana* leaves in which PhAvh202 or RFP was transiently expressed. The result showed that the lesion area was larger in the presence of PlAvh202 than that of RFP (Figure 3C, D). In addition, the biomass of *P. capsici* was also significantly higher in PlAvh202-expressing leaves compared with RFP (Figure 3E). These results suggested PlAvh202 could promote the infection of *P. capsici* in *N. benthamiana*.

### A C-terminal internal repeat (IR2) is required for PlAvh202-mediated ICD suppression

The analysis of amino acid sequence from SMART (http://smart.embl-heidelberg.de/) showed that PlAvh202 was composed of an N-terminal signal peptide (SP, 1-23 aa), a typical RXLR-EER motif (24-49 aa), and two C-terminal internal repeat motifs (IR1, 67-130 aa and IR2, 273-338 aa). Therefore, we constructed four truncated constructs of PlAvh202 without the SP to identify the functional region of PlAvh202 (Figure 4A). These four constructs (M1, M2, M3, and M4) were transiently expressed in *N. benthamiana*, as confirmed by western blotting (Figure 4B). As shown in Figure 4A, the truncation M1 containing only the RXLR-EER region lost its ability to suppress ICD, demonstrating that the RXLR-EER motif could not suppress ICD. In contrast, the truncation M2, which does not contain the RXLR region, retained the ability to suppress ICD, suggesting that IR1 and IR2 might be critical for ICD suppression. To further investigate the contribution of IR1 and IR2, we individually deleted IR1 or IR2 (M3 or M4) and tested their ability of ICD suppression. INF1-induced *N. benthamiana* cell death was observed in the leaves expressing M4 but not M3 version of PlAvh202, suggesting that IR2 plays an essential role in ICD suppression.

**Figure 4.**
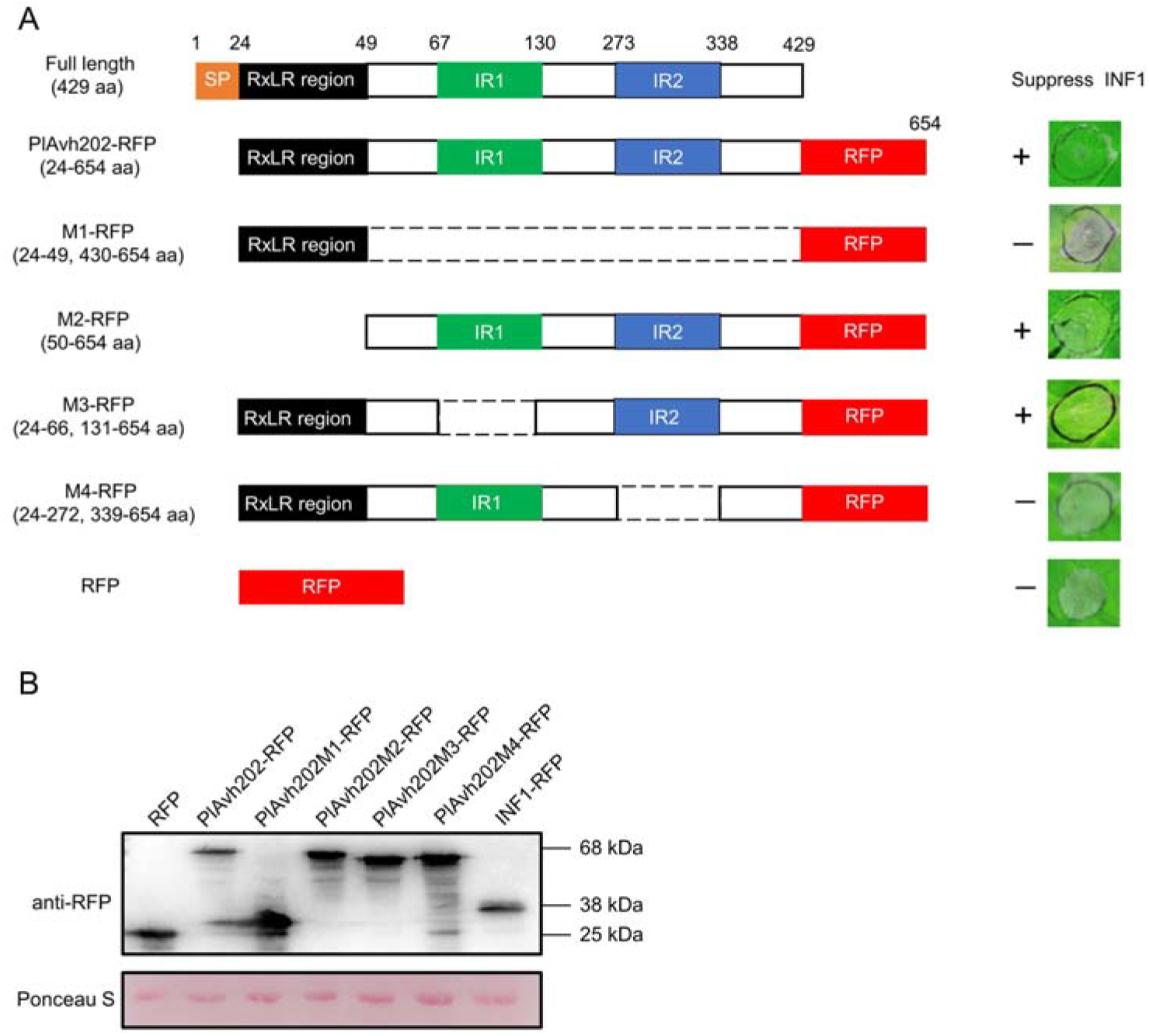
Internal repeat (IR2) is required for PlAvh202 to suppress ICD. **(A)** ICD suppression of PlAvh202 and its deletion mutants. Right panel is cell death symptoms in *N. benthamiana* leaves expressing PlAvh202 deletion mutants and INF1. Photographs were taken at 60 hpi, represents the ICD was not suppressed, “+” represents the ICD suppression. **(B)** Immunoblot analysis. INF1, RFP, PlAvh202 and its deletion mutants were detected by western blot using anti-RFP antibody. Total protein was stained by Ponceau S.

### Cytoplasmic localization of PlAvh202 is required for ICD suppression

To determinate the intracellular distribution of PlAvh202, PlAvh202-RFP was transiently co-expressed in *N. benthamiana* with a control GFP via agroinfiltration. PlAvh202-RFP was found at the periphery of cells and nucleus using laser confocal fluorescence microscopy (Figure 5A), suggesting that PlAvh202 might be located at plasma membrane (PM), cytoplasm, endoplasmic reticulum (ER) or nuclear envelope (NE). To further identify the specific subcellular location, PlAvh202 without SP was fused with an enhanced green fluorescent protein (eGFP) at its N-terminal and co-expressed with a PM maker AtPIP2A-RFP (Gonzalez et al., 2005) and an ER maker HDEL-mCherry (Fan et al., 2020), respectively. As shown in Figure 5A, GFP-PlAvh202 did not co-localize with AtPIP2A-RFP or HDEL-mCherry in a cell, indicating that PlAvh202 did not localize at PM or ER. Furthermore, we extracted membrane and cytosolic proteins of PlAvh202-RFP and AtPIP2A-RFP which was individually expressed *N. benthamiana*. Utilizing western blot, we could only detect PlAvh202-RFP in the cytosol fraction, while AtPIP2A-RFP was only detected in the membrane fraction (Figure 5B). This result excluded the possibility that PlAvh202 localized to cell membrane including PM, ER and NE. Thus, it demonstrates that PlAvh202 is a cytoplasmic protein.

**Figure 5.**
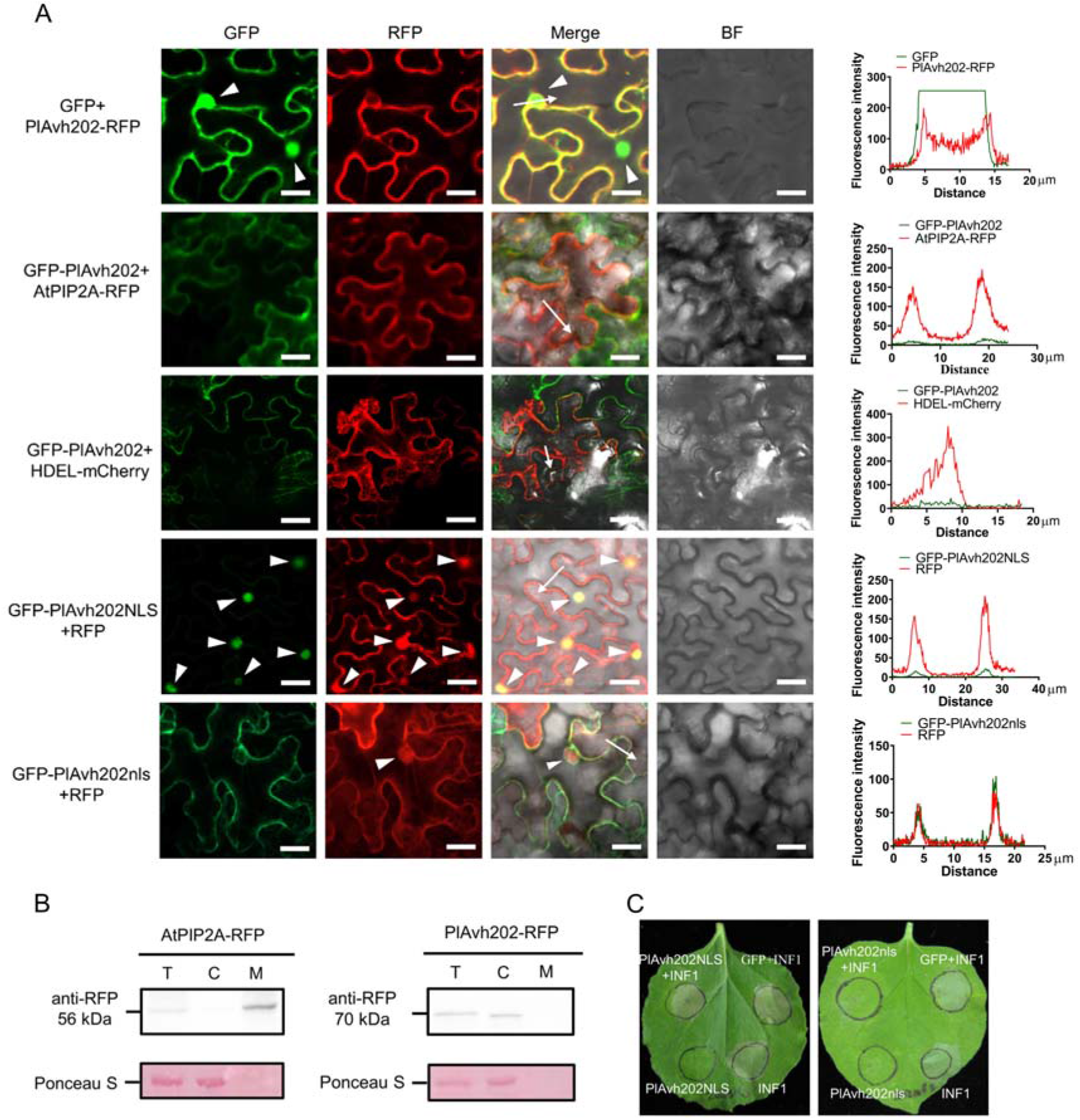
Cytoplasmic localization of PlAvh202 is required for ICD suppression. **(A)** The localization of PlAvh202. GFP-PlAvh202, GFP-PlAvh202^NLS^, GFP-PlAvh202^nls^ or PlAvh202-RFP was individually co-expressed with AtPIP2A-RFP (a plasma membrane maker), HDEL-mCherry (an endoplasmic reticulum maker), RFP or GFP in *N. benthamiana* leaf cells. Fluorescence was visualized by confocal microscopy at 36 hpi. Scale bars, 10 μm. A plot of the profile indicated by the white arrow shows GFP and RFP fluorescence signal at the region of interest, the white triangle indicated cell nucleus. Right panel is fluorescence intensity profiles of GFP and RFP. y axis, GFP or RFP relative fluorescence intensity; x axis, transect length (μm). **(B)** Western blot analysis of proteins from *N. benthamiana* leaves transiently expressing PlAvh202-RFP or AtPIP2A-RFP. T, total protein; C, cytosolic protein fraction; M, microsomal membrane protein fraction. **(C)** ICD suppression of PlAvh202 required cytoplasmic localization. *Agrobacterium* carrying *pBin::INF1::RFP* was infiltrated into *N. benthamiana* leaves in which GFP-PlAvh202^NLS^, GFP-PlAvh202^nls^ or GFP was expressed.

In addition, we also examined the subcellular location of M1, M2, M3, and M4, the result showed that M2, M3, and M4 were coincident with PlAvh202, but M1 had an extra nuclear localization (Figure S3A). In order to clarify if the nuclear localization of PlAvh202 has an impact on ICD suppression, we fused a nuclear localization signal (NLS) or mutated NLS (nls) to the C terminus of GFP-PlAvh202. The expression of GFP-PlAvh202^NLS^ and PlAvh202^nls^ proteins were confirmed by western blot (Figure S3B). Confocal imaging showed that GFP-PlAvh202^NLS^ was exclusively localized in nuclei, while GFP-PlAvh202^nls^ had the same subcellular localization pattern as GFP-PlAvh202 (Figure 5A). Moreover, PlAvh202^nls^ could suppress ICD as PlAvh202 in *N. benthamiana*, but PlAvh202^NLS^ lost the ability of ICD suppression (Figure 5C), These results suggested that the cytoplasmic localization of PlAvh202 is required for ICD suppression.

### PlAvh202 interacts with SAMS proteins of *P. litchii* in vivo and in vitro

To identify the host target of PlAvh202, total proteins were extracted from the *N. benthamiana* leaves transiently expressing GFP-tagged PlAvh202, and subject to immunoprecipitation (IP) for isolation of potential PlAvh202-interacting proteins. *N. benthamiana* expressing GFP served as a control. All purified proteins were analyzed utilizing liquid chromatography tandem-mass spectrometry (LC-MS/MS). Several potentially PlAvh202-associated proteins were detected, among which we identified a NbSAMS2-like protein (Figure S4A), and this protein was not detected in GFP-interacting proteins (Table S3). We successfully cloned the *NbSAMS2-like* and its homologs (*NbSAMS1* and *NbSAMS3*) based on *NbSAMS* sequences of NCBI (https://www.ncbi.nlm.nih.gov/) and Sol Genomics Network database (https://solgenomics.net/). GFP-NbSAMS1, GFP-NbSAMS2-like, GFP-NbSAMS3 or GFP was transiently co-expressed with PlAvh202-HA in *N. benthamiana* for Co-immunoprecipitation (Co-IP), PlAvh202-HA was co-immunoprecipitated with GFP-NbSAMS1, GFP-NbSAMS2-like, and GFP-NbSAMS3 but not GFP (Figure S5A). This result suggests that PlAvh202 interacts with all three NbSAMSs in vivo. Besides, a glutathione S-transferase (GST) pull-down assay of co-incubation between GST-tagged NbSAMSs protein and His-tagged PlAvh202 protein showed that His-tagged PlAvh202 could be pulled down by all three GST-tagged NbSAMSs but not GST control (Figure S5B), suggesting that PlAvh202 can interact with NbSAMS1, NbSAMS2-like, and NbSAMS3 in vitro.

To further validate the relationship between PlAvh202 and SAMS of litchi, four *LcSAMS* (*LcSAMS2*, *LcSAMS3*, *LcSAMS4*, and *LcSAMS5*) genes encoding the protein homologous to NbSAMS2-like were cloned from litchi cDNA and subsequently constructed into plasmid pCLuc, pBinGFP2 and pGEX-6P-1 for split luciferase complementation (SLC), Co-IP and pull-down. Both SLC and Co-IP assays confirmed that PlAvh202 could interact with each of four LcSAMSs in vivo (Figure 6A, B), and pull-down assays confirmed the interaction between PlAvh202 and all four LcSAMSs in vitro (Figure 6C). Plant SAMSs can be classified as Type I or II, and Type I is three times more abundant than Type II (Sekula et al., 2020). Although both LcSAMS2 and LcSAMS3 belong to Type I group (Figure S4B), LcSAMS3 was chosen for subsequent experiments because it is more similar to NbSAMS2-like in amino acid sequence (Figure S4C).

**Figure 6.**
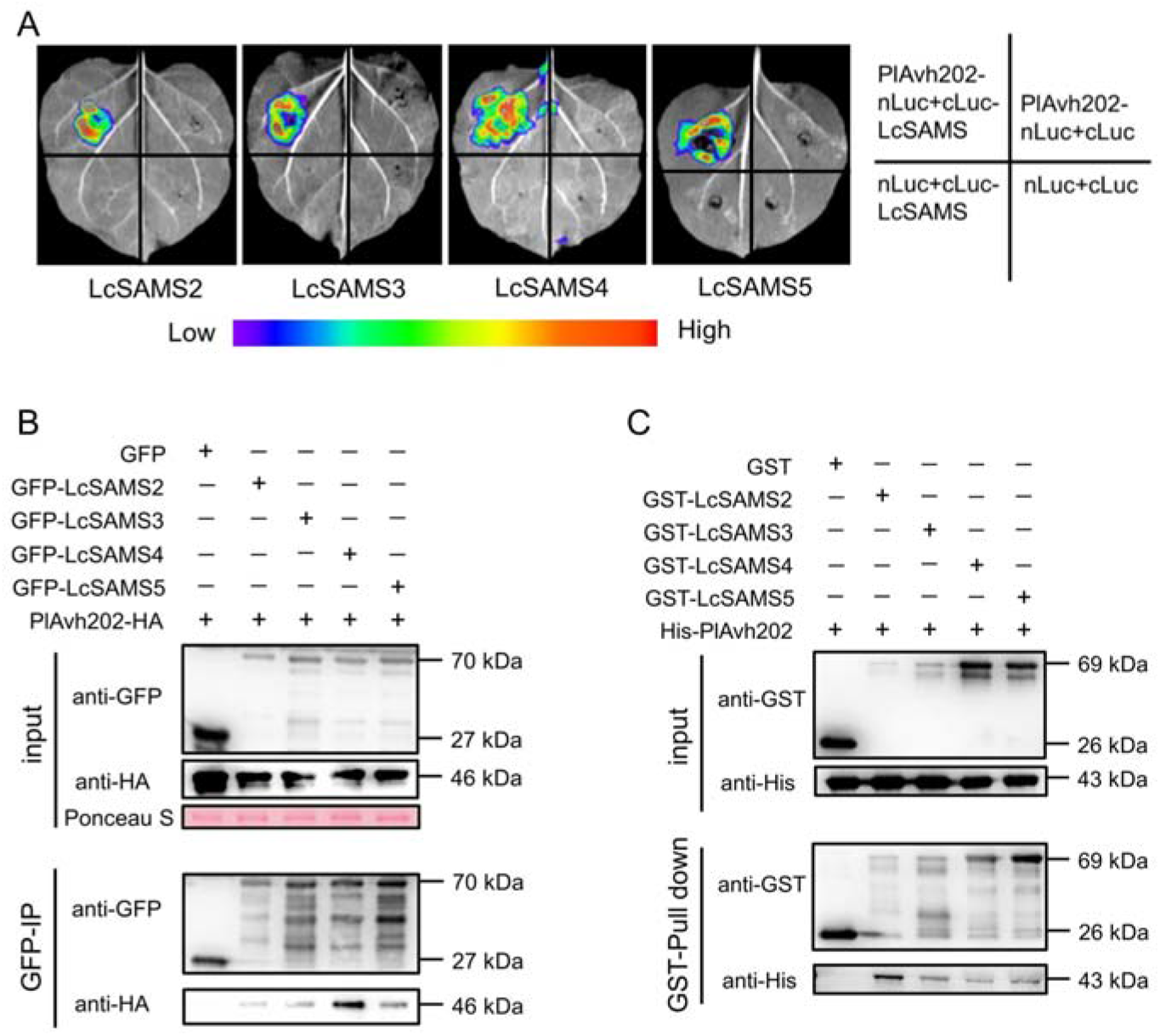
The interaction of PlAvh202 with LcSAMSs in vivo and in vitro. **(A)** Split luciferase complementation assays for the determination of interactions between PlAvh202 and LcSAMSs. PlAvh202-nLuc was co-expressed with cLuc-LcSAMS2, cLuc-LcSAMS3, cLuc-LcSAMS4 or cLuc-LcSAMS5 in *N. benthamiana* leaves through agroinfiltration. Fluorescence signal intensity was recorded at 3 dpa. **(B)** Co-immunoprecipitation of PlAvh202 by LcSAMSs. PlAvh202-HA was co-expressed with GFP-LcSAMS2, GFP-LcSAMS3, GFP-LcSAMS4 or GFP-LcSAMS5 in *N. benthamiana* leaves, and total protein was extracted at 60 hpi. Protein complexes were pulled down utilizing GFP-Trap beads and the captured proteins were detected by western blot using anti-HA antibody. Total protein was strained by Ponceau S. **(C)** In vitro pull-down assays of PlAvh202 by LcSAMSs. His-PlAvh202, GST-LcSAMS2, GST-LcSAMS3, GST-LcSAMS4 or GST-LcSAMS5 were expressed in *E. coli* and co-incubated as indicated in the input. Co-precipitation of His-PlAvh202 with GST-binding proteins were detected by western blot using anti-His antibody.

### SAMSs contribute to ICD and plant resistance against pathogens

To explore the role of SAMS in plant resistance, LcSAMS3 was transiently expressed in *N. benthamiana* for challenging with *P. capsici*. Expression of LcSAMS3 led to smaller lesions and less biomass of *P. capsici* than that caused by RFP (Figure 7A, S6A), suggesting that LcSAMS3 positively regulate plant resistance. In addition, *NbSAMSs* were silenced using virus-induced gene silencing (VIGS). As verified by RT-qPCR, the relative expression levels of three *NbSAMSs* declined by 70-90% in comparison with *GUS* control (Figure 7B). Although *NbSAMSs*-silenced plants showed a slower growth than control (Figure S6B), they could be used for subsequent experiments at 30 days post-agroinfiltration (dpa). PlAvh202 or RFP was transiently expressed in the control or *NbSAMS*s-silenced leaves, and *P. capsici* was inoculated in theses leaves after 24 h. The lesion area and biomass of *P. capsici* on PlAvh202-expressing leaves are the same as that on RFP-expressing leaves when *NbSAMS*s were silenced, whereas PlAvh202 still caused larger lesions and more biomass in the *GUS* control (Figure 7C, D). This result suggests that *NbSAMS*s are essential for PlAvh202 to promote *P. capsici* infection. Besides, enhanced susceptibility to *P. capsici* was also observed in *NbSAMS*s-silenced lines compared with *GUS* control (Figure 7C, D), highlighting the positive role of SAMS in regulating plant resistance to *P. capsici*. Moreover, a certain degree of ICD was diminished in *NbSAMS*s-silenced *N. benthamiana* although the ICD was still visible, indicating that *NbSAMS*s were involved in ICD (Figure 7C). In addition, RT-qPCR results showed that the relative expression levels of *NbRbohA* and *NbRbohB* in *NbSAMSs-silenced N. benthamiana* declined by 55% and 72% (Figure S6C), indicating that SAMS may contribute to ROS-based plant resistance and PCD. Taken together, these results support that PlAv202 diminishes ICD and enhances plant susceptibility via *NbSAMS*s.

**Figure 7.**
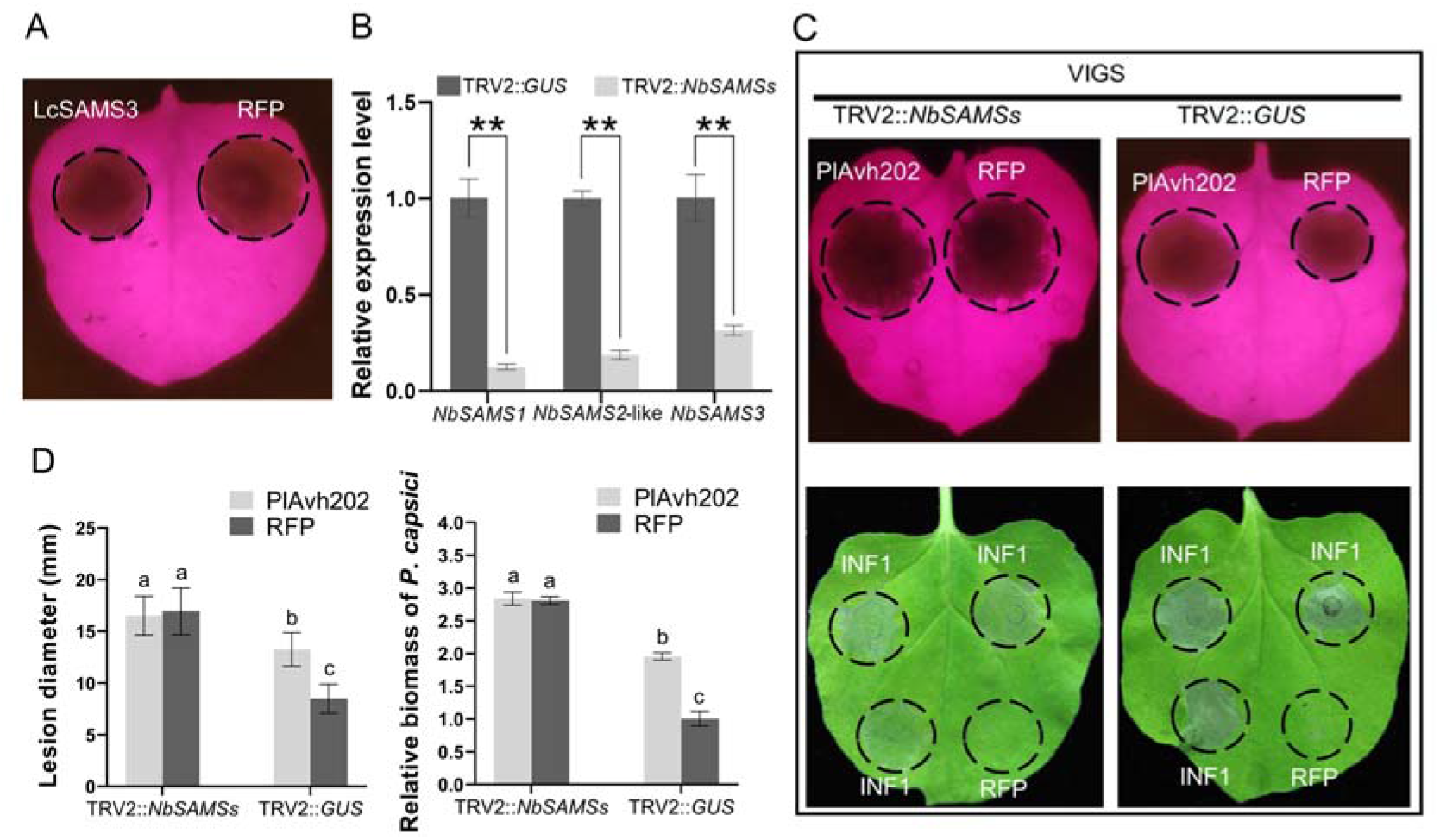
SAMSs positively regulate plant resistance. **(A)** Transient expression of LcSAMS3 enhanced the resistance of *N. benthamiana* to *P. capsici. P. capsici* was inoculated in PlAvh202 or RFP-expressing *N. benthamiana* leaves, and photographs were taken under UV light at 36 hpi. **(B)** Silencing efficiency of *NbSAMSs*. Relative expression levels of *NbSAMS1, NbSAMS2-like*, and *NbSAMS3* were measured by RT-qPCR. Data are the means ± SD of three independent biological replicates (Leaves sample of every experiment were from 3 plants), asterisks represent significant differences (\*\**P* < 0.01, Student’s *t* test). *NbEF1α* was used as the internal reference. **(C)** The silencing of *NbSAMSs* led to enhanced infection of *P. capsici* and diminished ICD. Top panel: *P. capsici* was inoculated in both sides of *NbSAMS*s-silenced *N. benthamiana* leaves expressing PlAvh202 and RFP, and photographs were taken at 36 hpi; bottom panel: INF1 or RFP was transiently expressed in *NbSAMS*s-silenced *N. benthamiana* leaves, and photographs were taken at 48 h post-agroinfiltration (hpa). TRV2::*GUS* was used as the control, black circles indicate lesion areas or cell death areas. **(D)** Lesion diameter and biomass of *P. capsici* in *NbSAMS*s-silenced *N. benthamiana* leaves. Different letters represent significant differences using the one-way ANOVA test followed by Tukey’s honestly significant difference (HSD) test (*P*< 0.05). Data are the means ± SD of three independent biological replicates (Leaves sample of every experiment were from at least 3 plants).

### Interaction with PlAvh202 leads to the destabilization of LcSAMSs

To determine which region of PlAvh202 is essential for interacting with LcSAMS3, GFP-tagged M1, M2, M3 and M4 were constructed to examine whether they could interact with LcSAMS3-HA using Co-IP. The results showed that LcSAMS3-HA was detected in the IP products of GFP-PlAvh202, GFP-PlAvh202M2, and GFP-PlAvh202M3, but not in that of GFP-PlAvh202M1, GFP-PlAvh202M4, and GFP (Figure 8A), suggesting that IR2 is required for the interaction between PlAvh202 and LcSAMS3. Interestingly, co-expression of PlAvh202 and LcSAMS3 appeared to reduce LcSAMS3 amounts in the Co-IP assay (Figure 8A). To further explore whether PlAvh202 alters LcSAMS3 stability, GFP, GFP-PlAvh202, and four mutants of GFP-PlAvh202 were co-expressed with LcSAMS3-HA in *N. benthamiana* to monitor the protein accumulation of LcSAMS3. According to the result of western blot, the accumulation of LcSAMS3-HA was reduced significantly in the presence of GFP-PlAvh202, GFP-PlAvh202M2 or GFP-PlAvh202M3 compared with GFP. However, the reduction of LcSAMS3 was not observed when LcSAMS3 was co-expressed with GFP-PlAvh202M1 or GFP-PlAvh202M4 (Figure 8B). This result suggests that PlAvh202 interacts with LcSAMS3 via IR2 motif, which can destabilize LcSAMS3.

**Figure 8.**
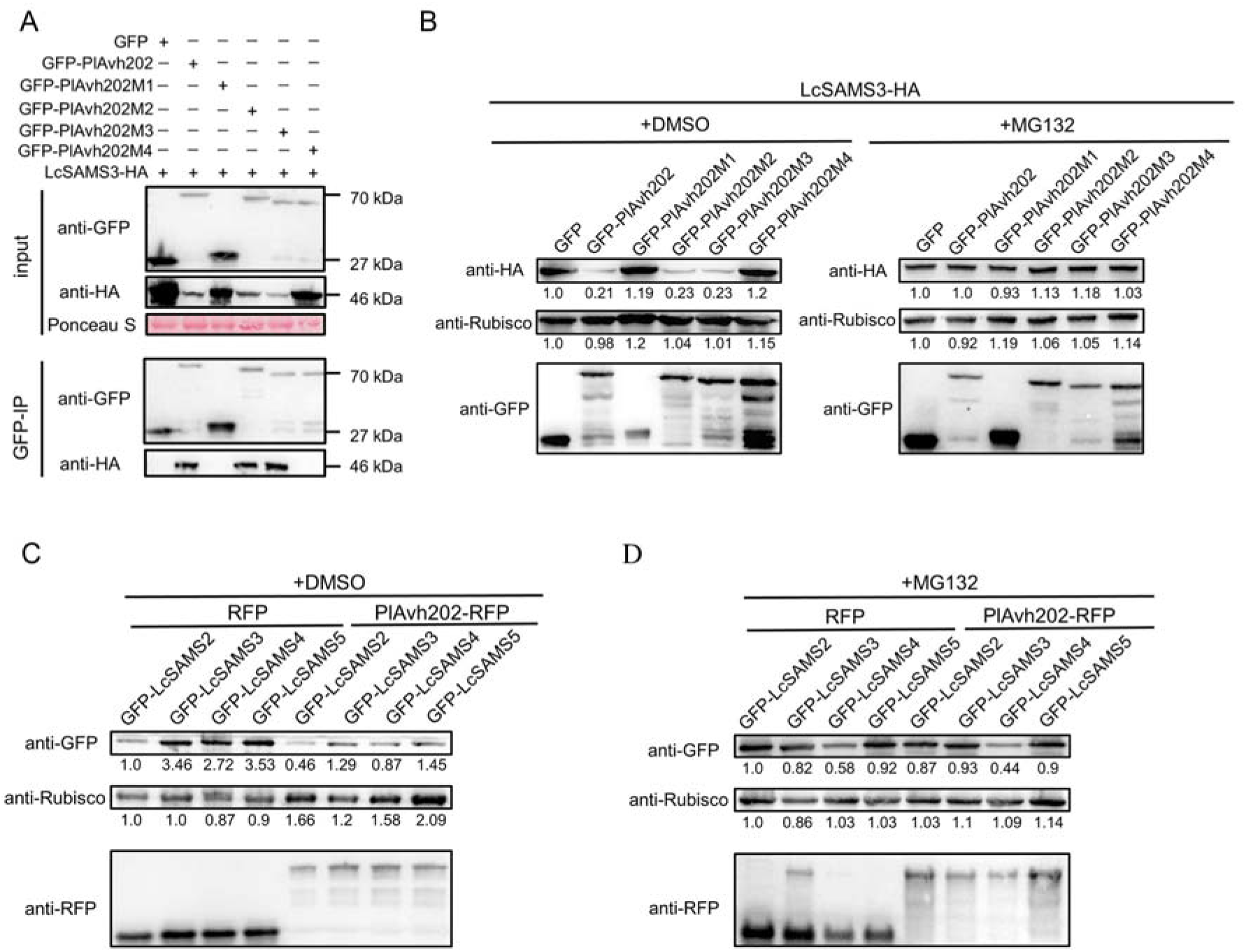
Association with PlAvh202 leads the degradation of LcSAMSs in a 26S proteasome-dependent manner. **(A)** Co-immunoprecipitation of LcSAMS3 by PlAvh202 and its mutants. LcSAMS3-HA was co-expressed with GFP-PlAvh202, GFP-PlAvh202M1, GFP-PlAvh202M2, GFP-PlAvh202M3 or GFP-PlAvh202M4 in *N. benthamiana* leaves, and total protein was extracted at 60 hpa. Protein complexes were pulled down utilizing GFP-Trap beads and the captured proteins were detected by western blot using anti-HA antibody. Total protein was strained by Ponceau S. **(B)** Protein stability of LcSAMS3 in the presence of PlAvh202 and its mutants. LcSAMS3-HA was co-expressed with GFP, GFP-PlAvh202, GFP-PlAvh202M1, GFP-PlAvh202M2, GFP-PlAvh202M3 or GFP-PlAvh202M4 in *N. benthamiana* leaves by agroinfiltration. These leaves were treated with dimethyl sulfoxide (0.5% DMSO, as a control) or MG132 (100 μM) at 48 hpa and total protein was extracted at 60 hpa for western blot. **(C, D)** Protein stability of LcSAMS2, LcSAMS3, LcSAMS4, and LcSAMS5 in the presence of PlAvh202 or RFP control. The procedures of leaf infiltration and treatment with DMSO **(C)** or MG132 **(D)** were described above. In **(B)-(D)**, relative protein abundance was indicated by the numbers below the blot, anti-Rubisco was used as the loading control.

In order to test whether the destabilization of LcSAMS3 by PlAvh202 depends on 26S proteasome, the assay of LcSAMS3 stability was repeated in the presence of MG132, which is an inhibitor of the 26S proteasome (Zhang et al., 2015). As shown in Figure 8B, the destabilization of LcSAMS3 by PlAvh202, PlAvh202M2, and PlAvh202M3 was inhibited by MG132 treatment. In addition, another three LcSAMSs (LcSAMS2, LcSAMS4 and LcSAMS5) were individually co-expressed with PlAvh202 in *N. benthamiana* to examine their protein stability. The result of western blot showed that PlAvh202 also destabilized these three LcSAMSs, and the destabilization could be inhibited by MG132 (Fiugre 8C, D). Taken together, these findings support that PlAvh202 can destabilize LcSAMSs in an interaction-dependent manner. Because IR2 is required for PlAvh202 to suppress ICD (Figure 4A) and destabilize SAMS (Figure 8A, B), and SAMS is involved in ICD (Figure 7C), we conclude that PlAvh202 targets and destabilizes plant SAMS to diminish ICD via IR2.

### PlAvh202-mediated ET suppression enhances plant susceptibility

To examine whether the stability of LcSAMS3 can impact ethylene production, LcSAMS3 was co-expressed with GFP-PlAvh202, GFP-PlAvh202M2, GFP-PlAvh202M3, GFP-PlAvh202M4, and GFP control, respectively, in *N. benthamiana* to measure ET concentrations by gas chromatography. As shown in Figure 9A, LcSAMS3 could induce ET production and the induced ET was significantly inhibited by GFP-PlAvh202, GFP-PlAvh202M2, GFP-PlAvh202M3 but not GFP-PlAvh202M4. Additionally, ET concentrations of MG132-treated plants were also measured. As excepted, ET concentrations inhibited by PlAvh202, GFP-PlAvh202M2, and GFP-PlAvh202M3 could be restored to GFP level in the presence of MG132 (Figure 9A). Given that PlAvh202, GFP-PlAvh202M2, and GFP-PlAvh202M3 but not GFP-PlAvh202M4 could target and destabilize LcSAMS3 (Figure 8A, B), it is a conclusion that PlAvh202 can destabilize LcSAMS3 via 26S proteasome to inhibit LcSAMS3-indeced ET production.

**Figure 9.**
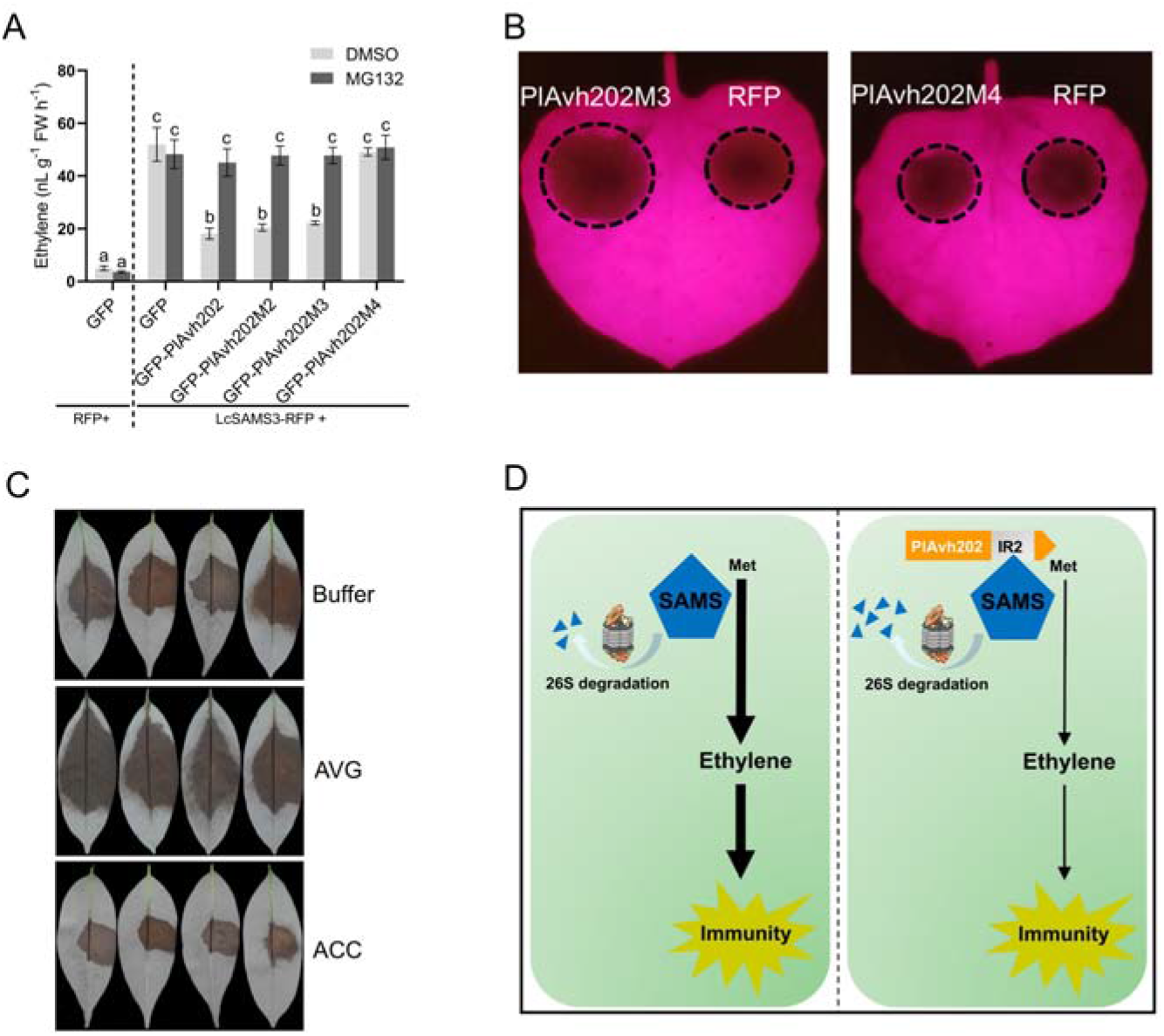
PlAvh202 enhances plant susceptibility by reducing LcSAMS-mediated ET production. **(A)** PlAvh202, PlAvh202M2, and PlAvh202M3 inhibit ET production induced by LcSAMS3 in *N. benthamiana* leaves. These leaves that LcSAMS3 was co-expressed with PlAvh202, PlAvh202M2, PlAvh202M3 or PlAvh202M4 were treated with DMSO or MG132 and weighed at 24 hpa, then the leaves were sealed in 10 mL glass vial to measure ET concentration using a gas chromatograph. Data are the means ± SD of three independent biological replicates (n≥6 leaves), different letters represent significant differences using the one-way ANOVA test followed by Tukey’s honestly significant difference (HSD) test (*P*< 0.05). **(B)** The reduced ET production led to enhanced susceptibility of plants to pathogens. RFP, PlAvh202M3 or PlAvh202M4 were transiently expressed in both sides of one leaf, and *P. capsici* was inoculated in these regions at 24 hpa. Photographs were taken under UV light at 36 hpi. **(C)** ET promotes litchi resistance to *P. litchii*. Litchi leaves were sprayed with 100 μM ACC, 50 μM AVG or 0.02% Silwet L-77 (as a control), and were maintained at high humidity for 3 h. These leaves were then inoculated 100 zoospores of *P. litchii* for infection analysis at 48 hpi. **(D)** A schematic diagram illustrating that PlAvh202 suppresses ET-mediated plant immunity by destabilizing SAMS. PlAvh202 interacts with plant SAMS via IR2 region, which leads to 26S proteasome-dependent degradation of SAMS. The destabilization of SAMS by PlAvh202 reduces ET production and ultimately results in the suppression of ET-mediated plant immunity.

To clarify the effect of reduced-ET production on PlAvh202 virulence, PlAvh202M3 or PlAvh202M4 was transiently expressed in *N. benthamiana* leaves to challenge with *P. capsici*. PlAvh202M3 was able to significantly promote *P. capsici* infection compared with RFP control, whereas PlAvh202M4 lost the ability to promote *P. capsici* infection (Figure 9B, S7A), suggesting that ET suppression is required for PlAvh202 virulence. To further determine the funtion of ET in *P. litchii* infection, we inoculated *P. litchii* on litchi leaves which were pretreated with 50 μM AVG (aminoethoxyvinylglycine, an ethylene biosynthesis inhibitor), 100 μM ACC (1-aminocyclopropane-1-carboxylic acid, a precursor of ethylene biosynthesis) or 0.02% Silwet L-77 buffer (as a control). Comparing with control, AVG-treated leaves displayed more severe symptoms of *P. litchii* infection, whereas ACC-treated leaves displayed less severe symptoms (Figure 9C, S7B), suggesting that ET production could promote litchi resistance to *P. litchii*. These results demonstrate that PlAvh202 promotes *P. litchii* infection by reducing ET production. Together, our findings suggest that PlAvh202 targets and destabilizes LcSAMSs to reduce ET production, which could enhance plant susceptibility (Figure 9D).

## Discussion

Effectors secreted from bacterial, fungal and oomycetous pathogens interact with host plants to interfere with plant PTI and ETI; thus, they are usually used as molecular probes to reveal immune mechanisms in the pathogen-plant interaction (Toruno et al., 2016). In our study, we found that an *P. litchii* RXLR effector, PlAvh202, had a strong ability to suppress INF1-triggered immune responses, including ICD, the expression of immune maker genes, and ROS accumulation, suggesting that PlAvh202 can suppress INF1-triggered PTI. In addition, PlAvh202 can also suppress Avr3a/R3a triggered cell death, suggesting that PlAvh202 has a potential ability to suppress Avr3a/R3a-triggered ETI. Although PTI and ETI are activated by two distinct mechanisms, they can cause many overlapping immune responses, including calcium influx, phosphorylation or dephosphorylation of mitogen-activated protein kinase (MAPK), ROS burst, and cell death (Adachi et al., 2015; Yuan et al., 2021), suggesting that some intersectant points in these two pathways. In fact, recent researches support that PTI and ETI are not independent with each other. Instead, they can produce some similar defense-related secondary metabolites and phytohormones to jointly enhance plants immune (Kadota et al., 2019; Yuan et al., 2021; Zhai et al., 2022). This opinion is consistent with our finding that PlAvh202 can suppress both INF1-triggered PTI and Avr3a/R3a-triggered ETI. Moreover, deletion of *PlAvh202* impaired the virulence of *P. litchii*. These results demonstrate that PlAvh202 has an important virulence function via suppressing plant immune. Therefore, PlAvh202 was used as a molecular probe to explore plant immune mechanism.

PlAvh202 contains two IRs, among which the C-terminal motif IR2 is required to suppress ICD and promote *P. capsici* infection. This result was coincident with recent reports that the IRs of RXLR effectors PsAvh23 and PlAvh142 were required for their virulence function (Kong et al., 2017; Situ et al., 2020). IR-containing proteins ubiquitously exist in all kingdoms of life and are responsible for binding diverse ligands, such as DNA, RNA, and proteins (Pawson and Nash, 2003; Bjorklund et al., 2006). Our study showed that IR2 is essential for PlAvh202 interacting with its plant target proteins, SAMSs, which is consistent with the protein-binding function of IRs. SAMSs make two contrary contributions to plant resistance. On the one hand, NbSAMS2 is required for the TGS (transcriptional gene silencing) and PTGS (post-transcriptional gene silencing) in *N. benthamiana* to resist the invading of *Cotton Leaf Curl Multan virus* (CLCuMuV) (Ismayil et al., 2018), suggesting that SAMS positively contributes to plant resistance. On the other hand, *Rice dwarf virus* (RDV)-encoded Pns11 protein could enhance the enzymatic activity of OsSAMS1 to significantly increase rice ET production, resulting in higher susceptibility of rice to RDV (Zhao et al., 2017). In our study, PlAvh202 could reduce LcSAMS-catalyzed ET production, leading to diminished plant resistance to *P. litchii*. This result is coincident with the role of SAMS in positively regulating plant resistance, whereas it may seem to be contrary to increased ET production resulting in the enhanced infection of RDV (Zhao et al., 2017).

ET plays a complicated role in plant-pathogen interactions, as it can positively or negatively regulated disease resistance (Broekaert et al., 2006; van Loon et al., 2006). For example, while ethylene-insensitive mutants of tobacco showed lower susceptibilities to *Peronospora parasitica*, they showed higher susceptibilities to *Colletotrichum destructivum* and *Fusarium oxysporum* (Chen et al., 2003; Geraats et al., 2003). Generally, the precise role of ET in plant immune responses depends on the pathogen type and environmental conditions (Yang et al., 2013; Washington et al., 2016). Although ET is usually produced by plants to restrict the invasion of *Phytophthora* pathogens (Nunez-Pastrana et al., 2011; Sugano et al., 2013; Shen et al., 2016; Shibata et al., 2016), the funtion of ET in plant resistance to *P. litchii* has not been reported. Our result that ET positively litchi resistance to *P. litchii* demonstrates that *P. litchii* is also susceptible to ET. Both flg22 and INF1 could induce ET production (Robert-Seilaniantz et al., 2011; Ohtsu et al., 2014), suggesting that ET contributes to plant PTI, which is consistent with our result that PlAvh202 inhibits INF1-induced PTI and ET production. Another RXLR effector, PsAvh238, could destabilize Type2 GmACS1 to reduce host plant ET production and facilitate *P. sojae* infection (Yang et al., 2019), suggesting that ET biosynthesis pathway is a significant target attacked by oomycete effectors to suppress plant immune. This research is coincident with our results that PlAvh202 impairs ET biosynthesis by destabilizing SAMSs to suppress plant immunity.

Plant PCD can be triggered by the crosstalk between some proteases with ROS, phytohormone (ET), and calcium ions (Huysmans et al., 2017; Sychta et al., 2021). *NbRbohA* and *NbRbohB* of *N. benthamiana* are mainly responsible for ROS production to enhance PCD and resist oomycete pathogens (Yoshioka et al., 2003), and therefore are involved in INF1 or Avr3a/R3a-triggered PCD. Our results showed that ICD and expression of *NbRbohA* and *NbRbohB* were reduced in *NbSAMS*s-silenced plants, suggesting that NbSAMSs may contribute to ICD through Rbohs-mediated ROS production. Moreover, previous study showed that INF1 could induce ET production (Ohtsu et al., 2014), and elevated ET level was essential for ROS burst and PCD (Qin and Lan, 2004; Liu et al., 2008; Yu et al., 2019). Given that SAMSs positively regulate ET production, it is a reasonable hypothesis that the destabilization of SAMSs by PlAvh202 reduces ET production, which compromises ROS accumulation and thus diminishes ICD. In addition, it remains unclear whether PlAvh202-mediated SAMSs destabilization is also involved in Avr3a/R3a-triggered PCD, which is worth further exploring in future.

It is notable that the ICD in *NbSAMS*s-silenced plants is still visible although the degree of cell death is diminished compared with control, suggesting that PlAvh202 suppresses ICD may not only depend on SAMSs-mediated ET pathway, but there may be other plant target(s) of PlAvh202 to regulate ICD. Indeed, some effectors could associate with multiple plant targets to interfere with host immunity. For example, AvrB from *P. syringae* interacts with RAR1, RIN4, and RIPK to suppress flg22-induced PTI (Shang et al., 2006; Lee et al., 2015). Another effector AvrPtoB of *P. syringa* targets FLS2, BAK1, and ubiquitin to suppress plant immunity (Abramovitch et al., 2006; Goehre et al., 2008). Therefore, identifying more potential targets of PlAvh202, which are involved in ICD or Avr3a/R3a-triggered PCD, is important to explore PTI-ETI crosstalk.

In summary, our study demonstrates, for the first time, that an oomycete RXLR effector targets plant SAMSs to manipulate host ET-based immunity, revealing a new mechanism of oomycetous pathogen-host plant interaction.

## Materials and methods

### Microbe and plant cultivation

*Peronophythora litchii* strain (SHS3) and *P. capsici* strain (PcLT263) were grown on CJA (carrot juice agar) medium in the dark at 25°C. *Escherichia coli* strains DH5α, JM109, BL21 and *Agrobacterium tumefaciens* (GV3101) were cultured on LB (Luria-Bertani) agar medium at 37 and 28°C, respectively. *N. benthamiana* plants were grown in greenhouse at 25°C with 16 hours light and 8 hours darkness.

### Plasmid construction

All primers in this study were listed in Table S2. *PlAvh202* and other *P. litchii* RXLR genes (without signal peptide-encoding sequence) were amplified from the cDNA of *P. litchii*, *NbSAMS* and *LcSAMS* genes were amplified from the cDNA of *N. benthamiana* and *litchi*. These amplified fragments were digested with *Sma*I and cloned into pBinRFP and pBinGFP2 using ClonExpress II One Step Cloning Kit (Vazyme) for transient expression of *N. benthamiana*. In addition, the amplified fragments of *PlAvh202* and *SAMS* were digested with *Bam*HI and *Eco*RI, respectively, and cloned into pET32a and pGEX-6P-1 for protein expression and purification in *E. coli*. The vectors pYF2.3G-RibosgRNA and pBluescript II KS were used for the deletion of *PlAvh202* by CRISPR/Cas9-mediated genome editing technology (Fang et al., 2017; Situ et al., 2020).

### Reverse transcription-quantitative PCR (RT-qPCR) assay

The total RNA of plant and microbe samples were extracted using an All-In-One DNA/RNA Mini-preps Kit (Bio Basic) following operation manual. gDNA buffer was added into total RNA to remove DNA impurity and then the purified RNA was used as a template to synthesized cDNA using a FastKing RT Kit (TIANGEN). The RT-qPCR assays were performed on qTOWER^3^ Real-Time PCR thermal cyclers (Analytik Jena) in 20 μl reactions that contained 6.4 μl deionized water, 10 μl SYBR Premix ExTaq II (Takara), 0.4 μM gene-specific primers and 20 ng cDNA. The specific reaction conditions are as follows: 95°C for 2 min, 40 cycles at 95°C for 30 s, and 60°C for 30 s, followed by a dissociation progress, 95°C for 15 s, 60°C for 1 min, and 95°C for15 s.

### Transient expression in *N. benthamiana* by *A. tumefaciens* infiltration

Recombinant plasmids were introduced into competent cells of *A. tumefaciens* strain GV3101 via heat shock. For infiltration, these transformed *A. tumefaciens* containing plant expression plasmids were incubated at 180 rpm, 28°C for 36 h. The bacterial cells were collected and washed three times with 10 mM MgCl_2_, 10 mM MES, pH 5.7, and 100 μM acetosyringone at 4000 rpm for 4 min. Resuspended *A. tumefaciens* was infiltrated into 5-6 week-old *N. benthamiana* leaves by a 1 ml needleless syringe at optical density (OD_600_) of 0.4-0.6.

### CRISPR/Cas9 mediated gene knockout of *PlAvh202*

The transformation of *P. litchii* was performed to delete *PlAvh202* as described previously (Fang et al., 2017; Situ et al., 2020). In brief, the plasmid pYF2.3G-Ribo-sgRNA and pBluescript II KS of *PlAvh202* were constructed and co-transformed with pYF2-PsNLS-hSpCas9 into protoplasts of strain SHS3 mediated by polyethylene glycol (PEG). The transformants were screened on CJA plate supplemented with 50 μg/mL G418. Then, these candidate transformants were subjected to extracted genomic DNA for PCR and sequencing.

For virulence analysis of transformants, 100 zoospores were inoculated on the tender leaves of litchi (Guiwei) in the dark at 25°C. Each transformant was inoculated at least 8 leaves. The lesion diameter was measured at 48 hpi.

### Diaminobenzidine (DAB) staining

Effectors were transiently expressed in *N. benthamiana* and INF1 was expressed in the same site by agroinfiltration after 24 h. The ROS induced by INF1 accumulated for 36 h and then these leaves of *N. benthamiana* were collected to be stained with 1 mg/mL DAB for 8 h in the dark. The leaves were boiled in absolute ethanol for 10 min to be for decolorization.

### *Phytophthora capsici* inoculation assay

Two equal area grown mediums of *P. capsici* were inoculated on two sides of *N. benthamiana* leaves, respectively, in which *A. tumefaciens* was infiltrated 24 hours ago. The leaves were photographed under UV light at 36 hpi and then collected to extracted DNA for biomass analysis by qPCR.

### Confocal microscopy

Fluorescent proteins were transiently expressed in *N. benthamiana* leaves using agroinfiltration, subcellular localizations were observed with LSM 7810 (Carl Zeiss) laser scanning microscope at 36 hours post-infiltration. The excitation wavelength of GFP and RFP was 488 and 561 nm, respectively.

### Protein extraction and western blots

*Nicotiana benthamiana* leaves were collected at 36 hours post-infiltration and ground in liquid nitrogen to be powder. About 100 mg plant powder and 0.1 mM protease inhibitor PMSF (no. ST507; Beyotime) was added into 600 μL extraction buffer (no. P0013B; Beyotime) to release total protein. Besides, a protein extraction kit (P0033; Beyotime) was used for membrane and cytosol fractions (Yu et al., 2012). According to the product standard protocol, about 200 mg plant powder was dissolved into 1 mL buffer A, vortexed 30 s, and then centrifuged at 700 g for 10 min at 4°C. Transferring supernatant to a new Eppendorf tube and centrifuged at 14000 g for 30 min at 4°C to precipitate membrane fraction, the supernatant (cytosol fraction) was collected in another new Eppendorf tube. The rest was centrifuged at 14000 g for 10 s at 4°C and removed supernatant as far as possible to reduce contamination of cytosol fractions. The precipitation was dissolved with 200 μL buffer B, which was vortexed for 5 s and placed on ice for 10 min. Vortex and ice-bath was repeated twice. Then the sample was centrifuged at 14000 g for 5 min at 4°C, the supernatant was membrane fraction. The protein sample was boiled at 100°C for 8 min with loading buffer and then separated on 12% sodium dodecyl sulfate–polyacrylamide gel electrophoresis (SDS-PAGE). After electrophoresis, the protein was treated as a standard protocol of western blot including membrane-transfer of polyvinylidene fluoride (PVDF), blocking with 5% nonfat dry milk, overnight incubation of monoclonal antibodies (Chromotek) on ice at a 1:5000 dilution, PBST washing (three times), incubation of goat anti-mouse horseradish peroxidase-conjugated secondary antibody (Dingguo) at a 1:5000 dilution and PBST washing (three times). Finally, the proteins were visualized using ECL reagents in the imaging system (Bio-Rad).

### Co-immunoprecipitation (Co-IP) and liquid chromatography-tandem mass spectrometry (LC-MS/MS) analysis

Genes were constructed into pBinGFP2 for transient expression of *N. benthamiana*. At 36 hpa, the protein was extracted from 1 g leaves using 2 mL extraction buffer (no. P0013; Beyotime) with 0.1 mM PMSF (no. ST507; Beyotime) and 0.1% (v/v) protease inhibitor cocktail (Sigma-Aldrich). The protein sample was centrifuged at 14000 g for 10 min at 4°C and then the supernatant was transferred in a new Eppendorf tube to incubate with 20 μl GFP-Trap-M beads (Chromotek) for 1 h at 4°C. After that, the beads were collected by a magnetic grate of DynaMag^TM^-2 (Invitrogen) and washed four times with wash buffer (10% glycerol, 1 mM EDTA, 150 mM NaCl, 25 mM Tris-Cl (pH 7.5), 0.5% Triton X-100 and 0.1% (v/v) protease inhibitor cocktail). For LC-MS/MS analysis, the beads were boiled with 40 μl SDT (4% (w/v) SDS, 100 mM Tris/HCl, 1 mM DTT (pH 7.6)) for 10 min and then sequenced by Beijing Genomics Institute (Shenzhen, China). For Co-IP, the beads were boiled with 40 μl protein loading buffer for 10 min, and then were used for western blotting.

### GST pull-down

The plasmids pGEX-6P-1, pGEX-6P-1-*SAMS* and *pET32a-PlAvh202* were expressed in *E. coli* strain BL21 to produce GST, GST-tagged SAMS and His-tagged PlAvh202 proteins in the condition of 18°C, 180 rpm and 0.1 mM isopropyl-β-D-thiogalactopyranoside (IPTG) for 10 h. These bacteria were pelleted and lysed using a sonic dismembrator (WIGGENS) in the cold lysis buffer (1×PBS (pH 7.4), 1% Triton X-100 and 0.1% (v/v) protease inhibitor cocktail) to release soluble proteins. GST and GST-tagged SAMS were incubated with 40 μl GST magnetic beads (Thermo Fisher Scientific) for 1 h at 4°C and then these beads were washed three times with lysis buffer. After that, the beads were incubated with His-tagged PlAvh202 1 h at 4°C and then washed three times and boiled for 10 min. The denatured proteins were used for western blotting.

### Split luciferase complementation (SLC) assay

The coding sequence of PlAvh202 (without SP) or LcSAMSs was constructed into pCAMBIA1300-nLUC or pCAMBIA1300-cLUC, and then PlAvh202-nLuc was co-expressed with cLuc-LcSAMS in *N. benthamiana* via *A. tumefaciens*-mediated transient expression (described above). Three days post-agroinfiltration, 0.3 mg/mL D-luciferin (Yeasen, China) was smeared onto infiltrated area. Luciferase signals were imaged using a low-light cooled charge-coupled device imaging system (NightSHADE LB 985 system, Berthold Technologies, Germany).

### Virus-induced gene silencing (VIGS) assays

The *A. tumefaciens* carrying *TRV2::NbSAMS*, TRV2::*GUS* or TRV1 was derived from preserved strains of our laboratory. Each of TRV2-harbored *A. tumefaciens* was mixed with TRV1 at a 1:1 ratio (each final OD_600_ was 0.25), TRV2::*GUS* was a control. Then they were co-infiltrated into 2-week-old *N. benthamiana* seedings. After 3 weeks, the upper leaves were used for RT-qPCR to analyze silencing efficiency and subsequent experiments.

### Ethylene quantification

*N. benthamiana* leaves treated by agroinfiltration 24 h ago were excised and weighed. Leaves were sealed in a 10 mL glass vial at 25°C for 6 h. 1 mL gas sample of head space was withdrawn and injected into the gas chromatography (Agilent 7890B) utilizing a gas-tight syringe to measure ethylene concentration. The column (Agilent, GS-Alumina; 50 m x 530 μm x 0 μm) was held at 50°C for 3 min. The temperature of sample entry and hydrogen flame ionization detector (FID) was 200°C and 300°C, respectively. The peak area from gas chromatography was used to calculate ethylene concentration according to the standard curve.

### Statistical analysis

Statistical analysis was performed using Prism version 8 (GraphPad, La Jolla, CA, USA). Student’s *t* test or one-way ANOVA with Tukey’s method was used to determine significance, and data are means ± SD of at least three independent experiments.

### Accession numbers

Sequence data from this article can be found in NCBI (https://www.ncbi.nlm.nih.gov/), SGN (https://solgenomics.net/) and SAP (http://www.sapindaceae.com/) databases. NCBI numbers: *NbSAMS1* (KX452091) and *NbSAMS3* (KX452093). SGN number: *NbSAMS2-like* (Niben101Scf01236g02016.1). SAP numbers: *LcSAMS2*(LITCHI014272.4), *LcSAMS3* (LITCHI014272.3), *LcSAMS4* (LITCHI024602.3), and *LcSAMS5* (LITCHI018653.1).

## Supplemental data

**Figure S1.** Expression profile of *PlAvh202*

**Figure S2.** Knock-out of *PlAvh202* by CRISPR/Cas9

**Figure S3.** Subcellar localization of PlAvh202 mutants

**Figure S4.** LC-MS/MS results and types of litchi SAMS

**Figure S5.** PlAvh202 interacts with NbSAMS1, NbSAMS2-like, and NbSAMS3

**Figure S6.** LcSAMS3 promotes *P. capsici* infection and silencing *NbSAMSs* reduces the relative expression levels of *NbRbohA* and *NbRbohB*

**Figure S7.** Ethylene (ET) contributes plant resistance

**Table S1.** RXLR effectors screening for ICD suppression

**Table S2.** Primers used in this study

**Table S3.** PlAvh202-associating proteins detected by co-immunoprecipitation (Co-IP) followed by liquid chromatography-tandem mass spectrometry (LC-MS/MS)

## Acknowledgments

We thank Professor Brett Tyler (Oregon State University, United States) for CRISPR/Cas9 vectors and Professor Yuanchao Wang (Nanjing Agricultural University, China) for pBinGFP2 and pBinRFP vectors. We are grateful to Professor Yizhen Deng (South China Agricultural University, China) for critical reading, helpful discussion and revision on the manuscript; to Professor Suomeng Dong (Nanjing Agricultural University, China) for the critical reading of the manuscript.

## Parsed Citations

**Abramovitch RB, Janjusevic R, Stebbins CE, Martin GB (2006) Type III effector AvrPtoB requires intrinsic E3 ubiquitin ligase activity to suppress plant cell death and immunity. Proceedings of the National Academy of Sciences of the United States of America 103: 2851-2856**

Google Scholar: <u>Author Only Title Only Author and Title</u>

**Abramovitch RB, Kim YJ, Chen SR, Dickman MB, Martin GB (2003) Pseudomonas type III effector AvrPtoB induces plant disease susceptibility by inhibition of host programmed cell death. Embo Journal 22: 60-69**

Google Scholar: <u>Author Only Title Only Author and Title</u>

**Adachi H, Nakano T, Miyagawa N, Ishihama N, Yoshioka M, Katou Y, Yaeno T, Shirasu K, Yoshioka H (2015) WRKY transcription factors phosphorylated by MAPK regulate a plant immune NADPH oxidase in Nicotiana benthamiana. Plant Cell 27: 2645-2663**

Google Scholar: <u>Author Only Title Only Author and Title</u>

**Armstrong MR, Whisson SC, Pritchard L, Bos JIB, Venter E, Avrova AO, Rehmany AP, Bohme U, Brooks K, Cherevach I, Hamlin N, White B, Frasers A, Lord A, Quail MA, Churcher C, Hall N, Berriman M, Huang S, Kamoun S, Beynon JL, Birch PRJ (2005) An ancestral oomycete locus contains late blight avirulence gene Avr3a, encoding a protein that is recognized in the host cytoplasm. Proceedings of the National Academy of Sciences of the United States of America 102: 7766-7771**

Google Scholar: <u>Author Only Title Only Author and Title</u>

**Axtell MJ, Staskawicz BJ (2003) Initiation of RPS2-specified disease resistance in Arabidopsis is coupled to the AvrRpt2-directed elimination of RIN4. Cell 112: 369-377**

Google Scholar: <u>Author Only Title Only Author and Title</u>

**Bjorklund AK, Ekman D, Elofsson A (2006) Expansion of protein domain repeats. PLoS Comput Biol 2: e114**

Google Scholar: <u>Author Only Title Only Author and Title</u>

**Broekaert WF, Delaure SL, De Bolle MF, Cammue BP (2006) The role of ethylene in host-pathogen interactions. Annu Rev Phytopathol 44: 393-416**

Google Scholar: <u>Author Only Title Only Author and Title</u>

**Casteel CL, De Alwis M, Bak A, Dong HL, Whitham SA, Jander G (2015) Disruption of ethylene responses by Turnip mosaic virus mediates suppression of plant defense against the green peach aphid vector. Plant Physiology 169: 209-218**

Google Scholar: <u>Author Only Title Only Author and Title</u>

**Chen N, Goodwin PH, Hsiang T (2003) The role of ethylene during the infection of Nicotiana tabacum by Colletotrichum destructivum Journal of Experimental Botany 54: 2449-2456**

Google Scholar: <u>Author Only Title Only Author and Title</u>

**DeFalco TA, Zipfel C (2021) Molecular mechanisms of early plant pattern-triggered immune signaling. Mol Cell 81: 3449-3467**

Google Scholar: <u>Author Only Title Only Author and Title</u>

**Ding P, Ding Y (2020) Stories of salicylic acid: a plant defense hormone. Trends Plant Sci 25: 549-565**

Google Scholar: <u>Author Only Title Only Author and Title</u>

**Dou D, Kale SD, Wang X, Chen Y, Wang Q, Wang X, Jiang RH, Arredondo FD, Anderson RG, Thakur PB, McDowell JM, Wang Y, Tyler BM (2008) Conserved C-terminal motifs required for avirulence and suppression of cell death by Phytophthora sojae effector Avr1b. Plant Cell 20: 1118-1133**

Google Scholar: <u>Author Only Title Only Author and Title</u>

**Du J, Verzaux E, Chaparro-Garcia A, Bijsterbosch G, Keizer LC, Zhou J, Liebrand TW, Xie C, Govers F, Robatzek S, van der Vossen EA, Jacobsen E, Visser RG, Kamoun S, Vleeshouwers VG (2015) Elicitin recognition confers enhanced resistance to Phytophthora infestans in potato. Nat Plants 1: 15034**

Google Scholar: <u>Author Only Title Only Author and Title</u>

**Dutra D, Agrawal N, Ghareeb H, Schirawski J (2020) Screening of secreted proteins of Sporisorium reilianum f. sp. zeae for cell death suppression in Nicotiana benthamiana. Front Plant Sci 11: 95**

Google Scholar: <u>Author Only Title Only Author and Title</u>

**Fabro G, Steinbrenner J, Coates M, Ishaque N, Baxter L, Studholme DJ, Korner E, Allen RL, Piquerez SJ, Rougon-Cardoso A, Greenshields D, Lei R, Badel JL, Caillaud MC, Sohn KH, Van den Ackerveken G, Parker JE, Beynon J, Jones JD (2011) Multiple candidate effectors from the oomycete pathogen Hyaloperonospora arabidopsidis suppress host plant immunity. PLoS Pathog 7: e1002348**

Google Scholar: <u>Author Only Title Only Author and Title</u>

**Fan XN, Che XR, Lai WZ, Wang SJ, Hu WT, Chen H, Zhao B, Tang M, Xie XA (2020) The auxin-inducible phosphate transporter AsPT5 mediates phosphate transport and is indispensable for arbuscule formation in Chinese milk vetch at moderately high phosphate supply. Environmental Microbiology 22: 2053-2079**

Google Scholar: <u>Author Only Title Only Author and Title</u>

**Fang Y, Cui L, Gu B, Arredondo F, Tyler BM (2017) Efficient genome editing in the oomycete Phytophthora sojae using CRISPR/Cas9. Current protocols in microbiology 44: 21A.21.21-21A.21.26**

Google Scholar: <u>Author Only Title Only Author and Title</u>

**Geraats BPJ, Bakker P, Lawrence CB, Achuo EĄ Hofte M, van Loon LC (2003) Ethylene-insensitive tobacco shows differentially altered susceptibility to different pathogens. Phytopathology 93: 813-821**

Google Scholar: <u>Author Only Title Only Author and Title</u>

**Goehre V, Spallek T, Haeweker H, Mersmann S, Mentzel T, Boller T, de Torres M, Mansfield JW, Robatzek S (2008) Plant pattern-recognition receptor FLS2 is directed for degradation by the bacterial ubiquitin ligase AvrPtoB. Current Biology 18: 1824-1832**

Google Scholar: <u>Author Only Title Only Author and Title</u>

**Gomez-Gomez L, Boller T (2000) FLS2: an LRR receptor-like kinase involved in the perception of the bacterial elicitor flagellin in Arabidopsis. Molecular cell 5: 1003-1011**

Google Scholar: <u>Author Only Title Only Author and Title</u>

**Gonzalez E, Solano R, Rubio V, Leyva A, Paz-Ares J (2005) PHOSPHATE TRANSPORTER TRAFFIC FACILITATOR1 is a plant-specific SEC12-related protein that enables the endoplasmic reticulum exit of a high-affinity phosphate transporter in Arabidopsis. Plant Cell 17: 3500-3512**

Google Scholar: <u>Author Only Title Only Author and Title</u>

**Huysmans M, Saul Lema A, Coll NS, Nowack MK (2017) Dying two deaths - programmed cell death regulation in development and disease. Current Opinion in Plant Biology 35: 37-44**

Google Scholar: <u>Author Only Title Only Author and Title</u>

**Ismayil A, Haxim Y, Wang YJ, Li HG, Qian LC, Han T, Chen TY, Jia Q, Liu AY, Zhu SB, Deng HT, Gorovits R, Hong YG, Hanley-Bowdoin L, Liu YL (2018) Cotton Leaf Curl Multan virus C4 protein suppresses both transcriptional and post-transcriptional gene silencing by interacting with SAM synthetase. Plos Pathogens 14**

Google Scholar: <u>Author Only Title Only Author and Title</u>

**Jiang RHY, Tripathy S, Govers F, Tyler BM (2008) RXLR effector reservoir in two Phytophthora species is dominated by a single rapidly evolving superfamily with more than 700 members. Proceedings of the National Academy of Sciences of the United States of America 105: 4874-4879**

Google Scholar: <u>Author Only Title Only Author and Title</u>

**Jones JD, Dangl JL (2006) The plant immune system. Nature 444: 323-329**

Google Scholar: <u>Author Only Title Only Author and Title</u>

**Kadota Y, Liebrand TWH, Goto Y, Sklenar J, Derbyshire P, Menke FLH, Torres MA, Molina A, Zipfel C, Coaker G, Shirasu K (2019) Quantitative phosphoproteomic analysis reveals common regulatory mechanisms between effector- and PAMP-triggered immunity in plants. New Phytologist 221: 2160-2175**

Google Scholar: <u>Author Only Title Only Author and Title</u>

**Kale SD, Gu B, Capelluto DG, Dou D, Feldman E, Rumore A, Arredondo FD, Hanlon R, Fudal I, Rouxel T, Lawrence CB, Shan W, Tyler BM (2010) External lipid PI3P mediates entry of eukaryotic pathogen effectors into plant and animal host cells. Cell 142: 284-295**

Google Scholar: <u>Author Only Title Only Author and Title</u>

**Kale SD, Tyler BM (2011) Entry of oomycete and fungal effectors into plant and animal host cells. Cell Microbiol 13: 1839-1848**

Google Scholar: <u>Author Only Title Only Author and Title</u>

**Kong L, Qiu X, Kang J, Wang Y, Chen H, Huang J, Qiu M, Zhao Y, Kong G, Ma Z, Wang Y, Ye W, Dong S, Ma W, Wang Y (2017) A Phytophthora effector manipulates host histone acetylation and reprograms defense gene expression to promote infection. Curr Biol 27: 981-991**

Google Scholar: <u>Author Only Title Only Author and Title</u>

**Lee D, Bourdais G, Yu G, Robatzek S, Coaker G (2015) Phosphorylation of the plant immune regulator RPM1-INTERACTING PROTEIN4 enhances plant plasma membrane H(+)-ATPase activity and inhibits flagellin-triggered immune responses in Arabidopsis. Plant Cell 27: 2042-2056**

Google Scholar: <u>Author Only Title Only Author and Title</u>

**Li Q, Ai G, Shen D, Zou F, Wang J, Bai T, Chen Y, Li S, Zhang M, Jing M, Dou D (2019) A Phytophthora capsici effector targets ACD11 binding partners that regulate ROS-mediated defense response in Arabidopsis. Mol Plant 12: 565-581**

Google Scholar: <u>Author Only Title Only Author and Title</u>

**Li W, Han Y, Tao F, Chong K (2011) Knockdown of SAMS genes encoding S-adenosyl-L-methionine synthetases causes methylation alterations of DNAs and histones and leads to late flowering in rice. J Plant Physiol 168: 1837-1843**

Google Scholar: <u>Author Only Title Only Author and Title</u>

**Liu H, Wang Y, Xu J, Su T, Liu G, Ren D (2008) Ethylene signaling is required for the acceleration of cell death induced by the activation of AtMEK5 in Arabidopsis. Cell Research 18: 422-432**

Google Scholar: <u>Author Only Title Only Author and Title</u>

**Liu T, Ye W, Ru Y, Yang X, Gu B, Tao K, Lu S, Dong S, Zheng X, Shan W, Wang Y, Dou D (2011) Two host cytoplasmic effectors are required for pathogenesis of Phytophthora sojae by suppression of host defenses. Plant Physiol 155: 490-501**

Google Scholar: <u>Author Only Title Only Author and Title</u>

**Liu Y, Lan X, Song S, Yin L, Dry IB, Qu J, Xiang J, Lu J (2018) In planta functional analysis and subcellular localization of the oomycete pathogen Plasmopara viticola candidate RXLR effector repertoire. Front Plant Sci 9: 286**

Google Scholar: <u>Author Only Title Only Author and Title</u>

**Murphy F, He Q, Armstrong M, Giuliani LM, Boevink PC, Zhang W, Tian Z, Birch PRJ, Gilroy EM (2018) The potato MAP3K StVIK is required for the Phytophthora infestans RXLR effector Pi17316 to promote disease. Plant Physiol 177: 398-410**

Google Scholar: <u>Author Only Title Only Author and Title</u>

**Nunez-Pastrana R, Fabiola Arcos-Ortega G, Armando Souza-Perera R, Alberto Sanchez-Borges C, Elena Nakazawa-Ueji Y, Javier Garcia-Villalobos F, Alberto Guzman-Antonio A, Juan Zuniga-Aguilar J (2011) Ethylene, but not salicylic acid or methyl jasmonate, induces a resistance response against Phytophthora capsici in Habanero pepper. European Journal of Plant Pathology 131: 669-683**

Google Scholar: <u>Author Only Title Only Author and Title</u>

**Ohtsu M, Shibata Y, Ojika M, Tamura K, Hara-Nishimura I, Mori H, Kawakita K, Takemoto D (2014) Nucleoporin 75 is involved in the ethylene-mediated production of phytoalexin for the resistance of Nicotiana benthamiana to Phytophthora infestans. Mol Plant Microbe Interact 27: 1318-1330**

Google Scholar: <u>Author Only Title Only Author and Title</u>

**Park C, Lee HY, Yoon GM (2021) The regulation of ACC synthase protein turnover: a rapid route for modulating plant development and stress responses. Current Opinion in Plant Biology 63**

Google Scholar: <u>Author Only Title Only Author and Title</u>

**Pawson T, Nash P (2003) Assembly of cell regulatory systems through protein interaction domains. Science 300: 445-452**

Google Scholar: <u>Author Only Title Only Author and Title</u>

**Pieterse CMJ, Van der Does D, Zamioudis C, Leon-Reyes A. Van Wees SCM (2012) Hormonal modulation of plant immunity. In R Schekman, ed, Annual Review of Cell and Developmental Biology, Vol 28, Vol 28, pp 489-521**

Google Scholar: <u>Author Only Title Only Author and Title</u>

**Qin WM, Lan WZ (2004) Fungal elicitor-induced cell death in Taxus chinensis suspension cells is mediated by ethylene and polyamines. Plant Science 166: 989-995**

Google Scholar: <u>Author Only Title Only Author and Title</u>

**Robert-Seilaniantz A, Grant M, Jones JD (2011) Hormone crosstalk in plant disease and defense: more than just jasmonate-salicylate antagonism. Annu Rev Phytopathol 49: 317-343**

Google Scholar: <u>Author Only Title Only Author and Title</u>

**Sekula B, Ruszkowski M, Dauter Z (2020) S-adenosylmethionine synthases in plants: Structural characterization of type I and II isoenzymes from Arabidopsis thaliana and Medicago truncatula. International Journal of Biological Macromolecules 151: 554-565**

Google Scholar: <u>Author Only Title Only Author and Title</u>

**Shang Y, Li X, Cui H, He P, Thilmony R, Chintamanani S, Zwiesier-Vollick J, Gopalan S, Tang X, Zhou J-M (2006) RAR1, a central player in plant immunity, is targeted by Pseudomonas syringae effector AvrB. Proceedings of the National Academy of Sciences of the United States of America 103: 19200-19205**

Google Scholar: <u>Author Only Title Only Author and Title</u>

**Shen D, Chai C, Ma L, Zhang M, Dou D (2016) Comparative RNA-Seq analysis of Nicotiana benthamiana in response to Phytophthora parasitica infection. Plant Growth Regulation 80: 59-67**

Google Scholar: <u>Author Only Title Only Author and Title</u>

**Shibata Y, Ojika M, Sugiyama A, Yazaki K, Jones DA, Kawakita K, Takemoto D (2016) The full-size ABCG transporters Nb-ABCG1 and Nb-ABCG2 function in pre-and postinvasion defense against Phytophthora infestans in Nicotiana benthamiana. Plant Cell 28: 1163-1181**

Google Scholar: <u>Author Only Title Only Author and Title</u>

**Situ J, Jiang L, Fan X, Yang W, Li W, Xi P, Deng Y, Kong G, Jiang Z (2020) An RXLR effector PlAvh142 from Peronophythora litchii triggers plant cell death and contributes to virulence. Mol Plant Pathol 21: 415-428**

Google Scholar: <u>Author Only Title Only Author and Title</u>

**Situ J, Jiang L, SHAO Y, Kong G, Xi P, Jiang Z (2020) Establishment of CRISPR /Cas9 genome editing system in Peronophythora litchii. Journal of Fungal Research 18: 181-188**

Google Scholar: <u>Author Only Title Only Author and Title</u>

**Sugano S, Sugimoto T, Takatsuji H, Jiang CJ (2013) Induction of resistance to Phytophthora sojae in soyabean (Glycine max) by salicylic acid and ethylene. Plant Pathology 62: 1048-1056**

Google Scholar: <u>Author Only Title Only Author and Title</u>

**Sychta K, Slomka A, Kuta E (2021) Insights into Plant Programmed Cell Death Induced by Heavy Metals-Discovering a Terra Incognita. Cells 10**

Google Scholar: <u>Author Only Title Only Author and Title</u>

**Toruno TY, Stergiopoulos I, Coaker G (2016) Plant-pathogen effectors: cellular probes interfering with plant defenses in spatial and temporal manners. Annu Rev Phytopathol 54: 419-441**

Google Scholar: <u>Author Only Title Only Author and Title</u>

**Turnbull D, Yang L, Naqvi S, Breen S, Welsh L, Stephens J, Morris J, Boevink PC, Hedley PE, Zhan J, Birch PRJ, Gilroy EM (2017) RXLR effector AVR2 up-regulates a brassinosteroid-responsive bHLH transcription factor to suppress immunity. Plant Physiol 174: 356-369**

Google Scholar: <u>Author Only Title Only Author and Title</u>

**van Loon LC, Geraats BP, Linthorst HJ (2006) Ethylene as a modulator of disease resistance in plants. Trends Plant Sci 11: 184-191**

Google Scholar: <u>Author Only Title Only Author and Title</u>

**Wang Q, Han C, Ferreira AO, Yu X, Ye W, Tripathy S, Kale SD, Gu B, Sheng Y, Sui Y, Wang X, Zhang Z, Cheng B, Dong S, Shan W, Zheng X, Dou D, Tyler BM, Wang Y (2011) Transcriptional programming and functional interactions within the Phytophthora sojae RXLR effector repertoire. Plant Cell 23: 2064-2086**

Google Scholar: <u>Author Only Title Only Author and Title</u>

**Washington EJ, Mukhtar MS, Finkel OM, Wan L, Banfield MJ, Kieber JJ, Dangl JL (2016) Pseudomonas syringae type III effector HopAF1 suppresses plant immunity by targeting methionine recycling to block ethylene induction. Proc Natl Acad Sci U S A 113: E3577-3586**

Google Scholar: <u>Author Only Title Only Author and Title</u>

**Wawra S, Belmonte R, Lobach L, Saraiva M, Willems A, van West P (2012) Secretion, delivery and function of oomycete effector proteins. Curr Opin Microbiol 15: 685-691**

Google Scholar: <u>Author Only Title Only Author and Title</u>

**Whisson SC, Boevink PC, Moleleki L, Avrova AO, Morales JG, Gilroy EM, Armstrong MR, Grouffaud S, van West P, Chapman S, Hein I, Toth IK, Pritchard L, Birch PR (2007) A translocation signal for delivery of oomycete effector proteins into host plant cells. Nature 450: 115-118**

Google Scholar: <u>Author Only Title Only Author and Title</u>

**Yang B, Wang YY, Guo BD, Jing MF, Zhou H, Li YF, Wang HN, Huang J, Wang Y, Ye WW, Dong SM, Wang YC (2019) The Phytophthora sojae RXLR effector Avh238 destabilizes soybean Type2 GmACSs to suppress ethylene biosynthesis and promote infection. New Phytologist 222: 425-437**

Google Scholar: <u>Author Only Title Only Author and Title</u>

**Yang DL, Yang Y, He Z (2013) Roles of plant hormones and their interplay in rice immunity. Mol Plant 6: 675-685**

Google Scholar: <u>Author Only Title Only Author and Title</u>

**Ye W, Wang Y, Shen D, Li D, Pu T, Jiang Z, Zhang Z, Zheng X, Tyler BM, Wang Y (2016) Sequencing of the litchi downy blight pathogen reveals it is a Phytophthora Species with downy mildew-like characteristics. Mol Plant Microbe Interact 29: 573-583**

Google Scholar: <u>Author Only Title Only Author and Title</u>

**Yoshioka H, Numata N, Nakajima K, Katou S, Kawakita K, Rowland O, Jones JD, Doke N (2003) Nicotiana benthamiana gp91phox homologs NbrbohA and NbrbohB participate in H2O2 accumulation and resistance to Phytophthora infestans. Plant Cell 15: 706-718**

Google Scholar: <u>Author Only Title Only Author and Title</u>

**Yu X, Tang J, Wang Q, Ye W, Tao K, Duan S, Lu C, Yang X, Dong S, Zheng X, Wang Y (2012) The RxLR effector Avh241 from Phytophthora sojae requires plasma membrane localization to induce plant cell death. New Phytol 196: 247-260**

Google Scholar: <u>Author Only Title Only Author and Title</u>

**Yu Y, Wang J, Li S, Kakan X, Zhou Y, Miao Y, Wang F, Qin H, Huang R (2019) Ascorbic acid integrates the antagonistic modulation of ethylene and abscisic acid in the accumulation of reactive oxygen species. Plant Physiology 179: 1861-1875**

Google Scholar: <u>Author Only Title Only Author and Title</u>

**Yuan M, Jiang Z, Bi G, Nomura K, Liu M, Wang Y, Cai B, Zhou J-M, He SY, Xin X-F (2021) Pattern-recognition receptors are required for NLR-mediated plant immunity. Nature**

Google Scholar: <u>Author Only Title Only Author and Title</u>

**Yuan M, Ngou BPM, Ding P, Xin XF (2021) PTI-ETI crosstalk: an integrative view of plant immunity. Curr Opin Plant Biol 62: 102030**

Google Scholar: <u>Author Only Title Only Author and Title</u>

**Zebell SG, Dong X (2015) Cell-cycle regulators and cell death in immunity. Cell Host Microbe 18: 402-407**

Google Scholar: <u>Author Only Title Only Author and Title</u>

**Zhai K, Liang D, Li H, Jiao F, Yan B, Liu J, Lei Z, Huang L, Gong X, Wang X, Miao J, Wang Y, Liu JY, Zhang L, Wang E, Deng Y, Wen CK, Guo H, Han B, He Z (2022) NLRs guard metabolism to coordinate pattern- and effector-triggered immunity. Nature 601: 245-251**

Google Scholar: <u>Author Only Title Only Author and Title</u>

**Zhang MX, Li Q, Liu TL, Liu L, Shen DY, Zhu Y, Liu PH, Zhou JM, Dou DL (2015) Two cytoplasmic effectors of Phytophthora sojae regulate plant cell death via interactions with plant catalases. Plant Physiology 167: 164-175**

Google Scholar: <u>Author Only Title Only Author and Title</u>

**Zhao S, Hong W, Wu J, Wang Y, Ji S, Zhu S, Wei C, Zhang J, Li Y (2017) A viral protein promotes host SAMS1 activity and ethylene production for the benefit of virus infection. Elife 6**

Google Scholar: <u>Author Only Title Only Author and Title</u>

**Zipfel C, Kunze G, Chinchilla D, Caniard A, Jones JD, Boller T, Felix G (2006) Perception of the bacterial PAMP EF-Tu by the receptor EFR restricts Agrobacterium-mediated transformation. Cell 125: 749-760**

Google Scholar: <u>Author Only Title Only Author and Title</u>

